# Latexin facilitates neointimal formation via promoting smooth muscle cell proliferation and macrophage migration

**DOI:** 10.1101/2024.10.03.616555

**Authors:** Kyosuke Kazama, Wennan Liu, Zhi-Fu Guo, Xiujuan Zhang, Chen Zhang, Ross Summer, Jianxin Sun

## Abstract

**Rationale:** Neointimal formation is the major cause of cardiovascular diseases such as atherosclerosis and restenosis. Although restenosis rates are significantly reduced by the development of drug-eluting, the stent may not be implanted at all atherosclerotic sites, and long-term dual anti-platelet therapy is needed after stent implantation. Thus, it is essential to elucidate the detailed mechanisms underlying neointimal formation for developing novel therapies. Latexin (LXN), a previously identified pro-inflammatory protein, is highly expressed in vasculature and its expression is regulated by shear stress. Whether it plays a role in vascular remodeling, however, remains unknown.

**Objective:** To determine whether LXN is involved in neointimal formation, and if so, to define the molecular mechanisms involved.

**Methods and Results:** We found that the expression of LXN was significantly increased in neointimal hyperplasia, as determined by western blot. Immunofluorescent staining indicated that increased LXN expression was predominantly localized in smooth muscle cells and macrophages. To determine the cell specific roles of LXN in neointimal formation after injury, we generated global, smooth muscle cell (SMC)-specific, endothelial specific-, and myeloid-specific LXN knockout (KO) mice. We found that global, SMC-, and myeloid-specific LXN deficiency markedly prevented neointimal hyperplasia in mice after carotid artery ligation, whereas LXN deficiency in endothelial cells had no effects. Mechanistically, we found that LXN deficiency in SMCs significantly attenuated SMC proliferation and migration, mainly through inhibiting the expression of platelet-derived growth factor (PDGF) receptors. Intriguely, LXN deficiency in macrophages inhibited monocyte chemoattractant protein-1 (MCP-1)-induced macrophage migration through inhibiting ERK phosphorylation.

**Conclusions:** In summary, we for the first time demonstrated that LXN is essentially involved in SMC proliferation and microphage migration. Specific inhibition of LXN signaling may provide a novel therapeutic strategy for the treatment of cardiovascular diseases, such as restenosis and atherosclerosis.

**Novelty and Significance:** *What is known?:*

- Latexin (LXN) is considered to be a novel pro-inflammatory protein and its expression is responsive to laminar shear stress.
- Vascular inflammation, SMC proliferation, and macrophage migration are critical events in neointimal hyperplasia.

*What new information does this article contribute?:*

- LXN is elevated in carotid artery ligation-induced neointimal hyperplasia in mice.
- SMC-specific LXN deficiency prevents neointimal hyperplasia through inhibiting SMC proliferation migration via attenuating platelet-derived growth factor receptor expression.
- Myeloid-specific LXN deficiency prevents neointimal hyperplasia through inhibiting monocyte chemoattractant protein-1-induced macrophage migration and extracellular signal-regulated kinase phosphorylation. Neointimal formation is the major cause of cardiovascular disease such as atherosclerosis and restenosis after stenting or balloon angioplasty. Because of the development of drug-eluting stents, restenosis rates have been significantly reduced; however, it is still essential to explore the detailed mechanisms underlying neointimal hyperplasia. Latexin (LXN) is considered a novel pro-inflammatory protein that is ubiquitously expressed in vascular and immune cells. We found that LXN is elevated in neointimal hyperplasia, and its deficiency in SMCs and macrophages, but not in ECs, markedly prevents neointimal formation after carotid artery ligation in mice. Mechanistically, LXN deficiency in SMCs prevents SMC proliferation and migration via attenuation of platelet-derived growth factor receptor expression. Further, myeloid-specific LXN deficiency inhibits monocyte chemoattractant protein-1-induced macrophage migration via attenuation of extracellular signal-regulated kinase phosphorylation. For the first time, the present study demonstrated that LXN is a crucial mediator implicated in SMC biology and macrophage migration.

## Introduction

Neointimal hyperplasia is the major cause of restenosis by stenting or balloon angioplasty for atherosclerotic lesions, a progressive vascular remodeling defined as narrowing the arterial lumen^1^. Neointimal hyperplasia develops with damage to the arterial wall, which causes increased growth factors and chemoattractants, leading to proliferation and migration of vascular smooth muscle cells (SMCs) and recruitment and infiltration of immune cells ^1^. In healthy blood vessels, the SMC phenotype is largely contractile ^2^, however, in damaged blood vessels, SMCs switch to a synthetic phenotype with increased cell proliferation and migration, which are critically involved in the development of cardiovascular diseases, such as atherosclerosis and restenosis ^2^. Recently, restenosis rates have been significantly reduced by the development of drug-eluting ^3^. However, since stent may not be implanted at all atherosclerotic sites and the long-term dual anti-platelet therapy is needed after stent implantation, it is still essential to elucidate the mechanisms underlying neointimal hyperplasia and develop novel therapeutic targets ^3^.

Latexin (LXN) was initially identified in the lateral neocortex of the developing rat brain and is also known as an endogenous carboxypeptidase inhibitor ^4^. LXN comprises 222 amino acids and is widely expressed in various cells and tissues ^4^. Previously, LXN has been identified as a negative regulator of hematopoietic stem cells through concerted mechanisms of decreased self-renewal and increased apoptosis ^5–8^. Further, LXN has been shown to potentially function as a putative tumor suppressor protein ^9^ ^10^. Interestingly, LXN is highly expressed in vasculature and its expression is significantly downregulated in response to laminar shear stress, the role of LXN in regulating vascular hemostasis, however, remains largely unknown.

In the present study, we for the first time defined the cell-specific roles of LXN in vascular remodeling. We found that LXN was significantly increased in carotid artery ligation-induced neointimal hyperplasia, and its deficiency in SMCs and macrophages strikingly prevented the development of neointimal hyperplasia in mice after vascular injury Mechanistically, LXN deficiency prevents SMC proliferation by inhibiting platelet-derived growth factor (PDGF) receptor expressions and monocyte chemoattractant protein-1 (MCP-1)-induced macrophage migration by inhibiting ERK phosphorylation.

## Methods

### Data Availability

The authors declare that all supporting data are available within the article and its online supplementary files.

### Mice and Genotyping

All the animals were maintained at the facilities of Thomas Jefferson University, where meets the requirements of the Guide for the Care and Use of Laboratory Animals published by the US National Institutes of Health (NIH). The use of animals was approved by the Institutional Animal Care and Use Committee (IACUC) at Thomas Jefferson University. All mice used in this study were on the pure C57BL/6 background. LXN KO mice were purchased from Riken BRC (#RBRC03456). LXN floxed (fl) mice were purchased from Cyagen. Wild type (#000664), SM22 Cre (#004746), and Lyz2 Cre (#004781) mice were purchased from Jackson Laboratories (Bar Harbor, ME). VE-cadherin (Cdh5) Cre/ERT2 mice were obtained from Prof. Ralf H. Adams (The Max Planck Institute for Molecular Biomedicine, Göttingen, Germany). LXN^fl/fl^ mice were bred with SM22^Cre+/-^, Lyz2^Cre+/-^, or VE-cadherin Cre/ERT2^+/-^ to create SMC-specific (LXN^fl/fl^, SM22^Cre+/-^: LXN-sKO), myeloid-specific (LXN^fl/fl^, Lyz2^Cre+/-^: LXN-mKO), or endothelial cell (EC)-specific (LXN^fl/fl^, VE-cadherin Cre/ERT2^+/-^: LXN-eKO) LXN-KO mice, respectively. EC-specific LXN deficient was induced by intraperitoneal (i.p.) injection of tamoxifen (MP Biomedicals, S5960) in corn oil at a dose of 75 mg/kg body weight for five consecutive days, and mice were allowed to recover at least for one week before being used for experiments. Both sexes were used for all studies except endothelial studies. Genotyping was performed by polymerase chain reaction (PCR) at 2-3 weeks old with tail biopsies using primer pairs listed in Table 1 ^11^. All mice were housed under standard light conditions (12-hour light/12-hour dark cycle) and allowed free access to standard normal diet and water.

**Table 1:**
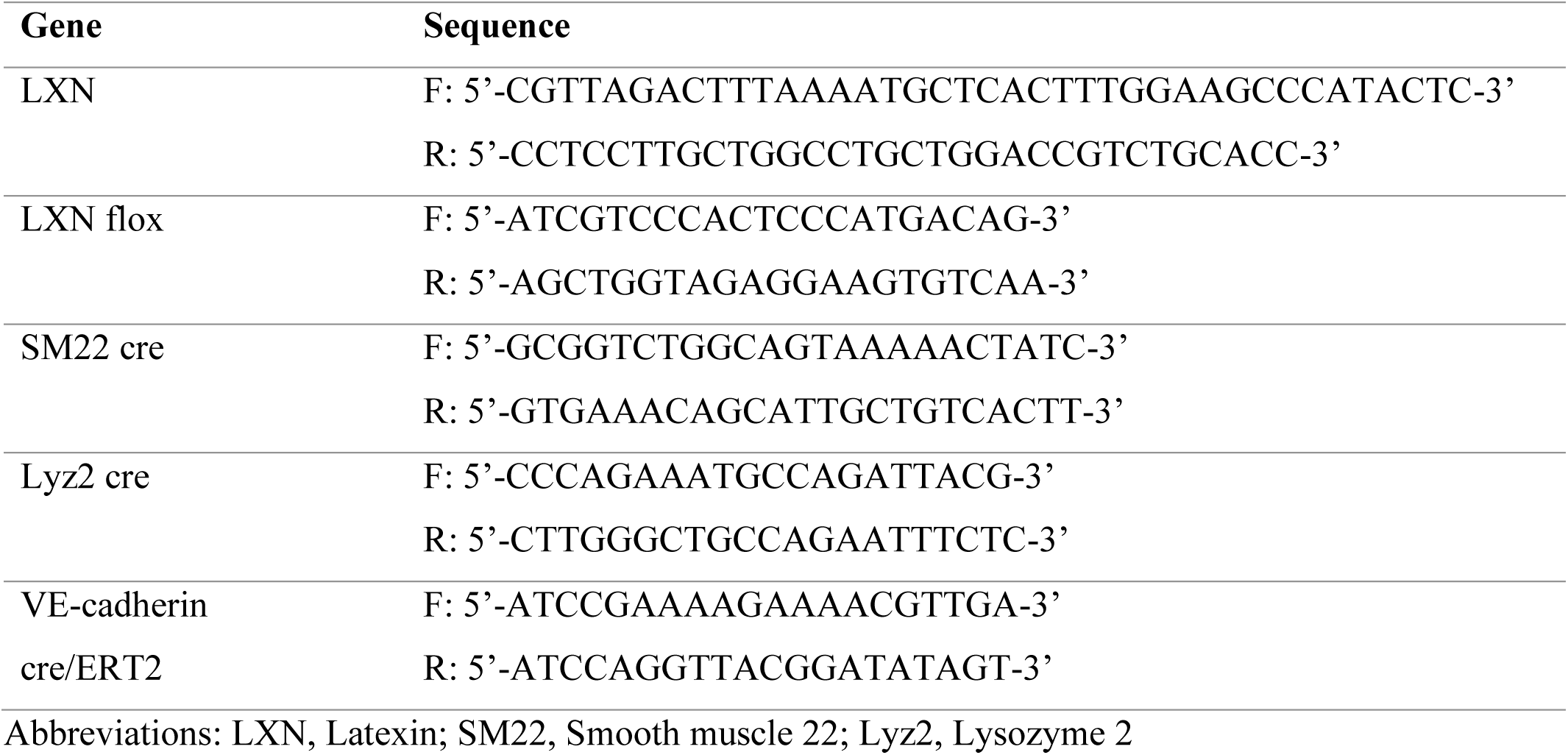
Primer pairs for genotyping PCR.

### Carotid artery ligation-induced neointimal hyperplasia

The carotid artery ligation was performed as previously described ^12^. Briefly, 2-4-month-old mice were anesthetized with 1-2% isoflurane. The skin was shaved and disinfected with iodine or ethanol. A 1.0-1.5 cm skin incision was made over the throat. Ligation surgery was performed on the left common carotid artery with a 6.0 silk suture, and the right common artery was used for sham control. The skin incision was closed with a 4.0 catgut suture. Buprenorphine (0.1-0.6 mg/kg, subcutaneous) was applied to reduce the pain before and after surgery as needed. 4-weeks after the surgery, mice were euthanized by exsanguination and perfusion under anesthesia, and carotid arteries were harvested and used for experiments.

### Primary cell isolation

Mouse SMCs were isolated from thoracic aortas as previously described ^13^. Briefly, three mice were used to isolate a sufficient number of SMCs. 3-4-week-old mice are euthanized by CO_2_ inhalation. Thoracic aortas were isolated under the dissection microscopy and placed into the P100 petri dish with calcium and magnesium-contained ice-cold Hanks’ Balanced Salt Solution (HBSS, 24020117, Gibco). Aortas were transferred into the 6-well plate and then incubated with 1 mg/ml collagenase II (Worthington Biochemical, LS004174) and 1 mg/ml soybean trypsin inhibitor (Worthington Biochemical, LS003570) in HBSS at 37 °C for 20 min. After removing adventitia and ECs, aortas were then cut into 1-2 mm pieces and incubated with 1 mg/ml collagenase II, 1 mg/ml soybean trypsin inhibitor, and 24.8 mg/ml elastase in HBSS at 37 °C for 30 min. Digested aortas were transferred into another 6-well plate and incubated in 20% Fetal bovine serum (FBS, 26140079, Gibco)-contained Dulbecco’s Modified Eagle’s Medium (DMEM)/F12 medium (10-090-CMR, Corning) supplemented with 1% penicillin/streptomycin (P/S, 10378016, Gibco) at 37 °C in a 5% CO_2_ humidified incubator for 1 week. Only low passage (3-5) of SMCs were used in this study.

Mouse peritoneal macrophages (PMs) were obtained from the peritoneal cavity as previously reported ^14^. Briefly, 2-3-month-old mice were euthanized, and skin and fur were removed from the peritoneum. 3 mL of phosphate buffered saline (PBS) with 1 mL of air was then injected into the abdominal cavity. 1 min after massaging, peritoneal cells in PBS were harvested and transferred into the 15 ml conical tube with 5 mL of serum-free Roswell Park Memorial Institute (RPMI) 1640 medium (10-043-CVR, Corning). After washing multiple times, peritoneal cells were incubated in 5% FBS-contained RPMI 1640 for 3 hours at 37 °C in a 5% CO_2_ humidified incubator. Non-adherent cells were removed by washing with PBS, and adherent cells were used as PMs in this study.

Bone marrow-derived macrophages (BMDMs) were obtained as previously reported ^14^. Briefly, 2-3-month-old mice were euthanized, and hind limb long bones (femurs and tibiae) were collected. Muscle or connective tissue was removed using forceps, scissors, and kimwipes. Push an 18 G needle through the bottom of a 0.5 ml tube and place the long bones into the tube. Nest the 0.5 ml tube in a 1.5 ml tube and then centrifuge at 10,000 g for 15 sec. Cells were incubated with 10 ml of red blood cell lysis buffer (Gibco, A10492) for 30 min and then filtered through a 70 μm Nylon cell strainer (Corning, 431751). Cells were centrifuged, dissociated with 10% FBS-contained RPMI 1640 medium, and incubated in a P100 petri dish at 37 °C. 4 h after the incubation, supernatants were collected and incubated with 10% FBS-contained RPMI 1640 medium supplemented with 10 ng/ml Macrophage Colony-Stimulating Factor (M-CSF, Sigma-Aldrich, M9170) for 5 days at 37 °C to differentiate into BMDMs.

ECs were obtained as previously reported ^15^. Briefly, lung lobes were isolated and washed in ice-cold HBSS. Lung tissues were minced and incubated in 15 mL of conditional medium containing 0.2 mg/mL type I collagenase (Worthington Biochemical Corporation) for 30 min at 37 ℃ with agitation. The digest was homogenized by passing multiple times through a 20-gauge needle with a 10 mL syringe, filtered through a 70 μm cell strainer, and then sorted with rat anti-mouse CD31 antibody (BD Biosciences)-conjugated with anti-rat IgG dynabeads (Thermo Fisher Scientific) using magnetic separation rack (New England Biolabs, Ipswich, MA). The beads/cell pellets were washed five times using the magnetic separator (Thermo Fisher Scientific) and used for experiments.

### Rat SMC culture and shRNA lentiviral infection

Rat aortic SMCs (Lonza, #R-ASM-580) were cultured in 5% FBS-contained DMEM (10-013-CV, Corning). Lentiviral pLKO-based short hairpin RNA (shRNA) oligonucleotides for rat LXN (target sequence: GGTATTCAAGGTGCAGACG) or non-targeting control (target sequence: GCGCGCTTTGTAGGATTCG) were transfected with packaging plasmids psPAX2 and pMD2.G to HEK293T cells using polyethylenimine (PEI, 1 mg/ml, #23966, Polysciences). Viral supernatant was harvested at 72 hrs post-transfection, and filtered through a 0.45 µm filter before adding to recipient SMCs, and supplemented with 10 µg/ml polybrene (Sigma-Aldrich, TR-1003). 24 hours after infection, the virus-containing media were removed and replaced with fresh media. Knockdown efficiency was detected 3 days after the virus infection.

### Ex vivo explant assay

Ex vivo mouse aortic explant assay was performed as previously reported ^16^. Aortas were isolated from LXN-gWT or -gKO mice, cleaned of perivascular adipose tissue and ECs, and cut into 2 mm segments. The segments were placed with the luminal side down and attach to the surface of a 6-well plate. Aortic segments were cultured with 20% FBS-contained DMEM medium. Outgrowth SMCs were counted with ImageJ software.

### Histology and Immuno-histochemistry (IHC)

Paraffin-embedded or frozen sections were used for histology and immune-histochemistry (IHC)^11^. For paraffin-embedded sections, harvested carotid arteries were fixed in 4% paraformaldehyde (PFA; Thermo Scientific, #9311) in PBS solution for 2 hours at room temperature. Tissues were then dehydrated, paraffinized, embedded, and sectioned using a microtome. For frozen sections, unfixed samples were embedded in Optimum Cutting Temperature (OCT, Sakura Finetek) compound, frozen, and sectioned using a cryostat. Serial sections were cut to 5 μm thickness onto Superfrost plus slides (Fisher Scientific, #12-550-15). Paraffin-embedded sections were deparaffinized and frozen sections were air-dried, and used for histology or IHC.

Hematoxylin (Poly Scientific, #72711) and Eosin (Poly Scientific, #71311) (H&E) staining were performed in accordance with the manufacturer’s instructions. IHC staining was performed as previously reported ^17^. Briefly, sections were washed three times with cold 0.1% Triton in PBS and then incubated with primary antibodies at 4 °C overnight after being blocked with protein blocking solution (DAKO) for 5-10 min at room temperature. Then, sections were washed three times with 0.1% Triton in PBS and incubated with HRP- or fluorescent-conjugated secondary antibodies for 1 hour at room temperature. Immune complexes with HRP-conjugated secondary antibodies were visualized with a diaminobenzidine (DAB) substrate kit (Abcam, ab64238). After dehydration, sections were rinsed three times with 0.1% Triton PBS and then mounted with Cytoseal XYL (Thermo Fisher Scientific, #8312-4). Primary and secondary antibodies used for IHC are listed in Table 2. Nuclei were stained with hematoxylin or 4’, 6-Diamidino-2-Phenylindole (DAPI; Thermo Fisher Scientific, D1306). Staining was observed under a Nikon eclipse 80i light microscope or a Nikon eclipse Ti2 confocal microscope. Quantification of the positive stained area was measured using ImageJ software provided by the NIH.

**Table 2:** Primary and secondary antibodies used for IHC.

| Antigen | Species | Dilution | Cat# | Source |
| --- | --- | --- | --- | --- |
| LXN | Rabbit | 1:1000 | ab154744 | Abcam |
| α-SMA | Mouse | 1:100 | sc-32251 | Santa Cruz |
| PCNA | Mouse | 1:100 | sc-56 | Santa Cruz |
| CD68 | Mouse | 1:100 | 66231 | Proteintech |
| Rabbit IgG (HRP-linked) | Goat | 1:1000 | 7074 | Cell signaling |
| Mouse IgG (HRP-linked) | Horse | 1:1000 | 7076 | Cell signaling |
| Rabbit IgG (Alexa Fluor 555) | Goat | 1:1000 | A21428 | Invitrogen |
| Mouse IgG (Alexa Fluor 488) | Donkey | 1:1000 | A21202 | Invitrogen |

Abbreviations: LXN, Latexin; α-SMA, alpha-smooth muscle actin; PCNA, Proliferating cell nuclear antigen; IgG, Immunoglobulin; HRP, Horseradish peroxidase

### Protein extraction

Isolated aortas, carotid arteries, or cultured cells were washed with cold PBS and then lysed with RIPA buffer [25 mM Tris-HCl, pH 7.6; 150 mM NaCl; 1% NP-40; 1% sodium deoxycholate; 0.1% sodium dodecyl sulfate (SDS)] supplemented with protease and phosphatase inhibitors ^12^. After incubating at 4 °C for 60 min, insoluble materials were removed by centrifugation. Protein concentration was determined using a bicinchoninic acid (BCA) protein assay kit (Pierce, #23225) in accordance with the manufacturer’s instructions.

### Western blot analysis

Western blot (WB) was performed as previously described ^12^. Briefly, equal amounts of protein (10-20 μg) were subjected to SDS-polyacrylamide gel (8-15%) electrophoresis and transferred to nitrocellulose (Bio-Rad, #1620115) membranes. After blocking with 3% bovine serum albumin (BSA; Fisher bioreagents, #BP9703) for 60 min at room temperature with gentle agitation, membranes were incubated with primary antibodies at 4 °C overnight, and membrane-bounded antibodies were visualized using fluorescent-conjugated secondary antibodies. Primary and secondary antibodies used for WB are listed in Table 3. Membranes were scanned using an Odyssey Imaging System (LI-COR). Equal loading of protein was confirmed by measuring β-actin or GAPDH expression. The results were analyzed using an Image Studio Lite Software (LI-COR).

**Table 3:** Primary and secondary antibodies used for WB.

| Antigen | Species | Dilution | Cat# | Source |
| --- | --- | --- | --- | --- |
| LXN | Rabbit | 1:1000 | ab154744 | Abcam |
| PCNA | Mouse | 1:1000 | sc-56 | Santa Cruz |
| Phospho-ERK | Rabbit | 1:2000 | 4370 | Cell signaling |
| Total-ERK | Mouse | 1:2000 | 4696 | Cell signaling |
| Phospho-NF-κB (Ser536) | Rabbit | 1:1000 | 3033 | Cell signaling |
| Total-NF-κB | Mouse | 1:1000 | sc-8008 | Santa Cruz |
| Phospho-Akt (Thr308) | Rabbit | 1:1000 | 9018 | Cell signaling |
| Total-Akt | Rabbit | 1:1000 | 9272 | Cell signaling |
| Phospho-p38 | Rabbit | 1:1000 | 4511 | Cell signaling |
| Total-p38 | Mouse | 1:1000 | sc-271120 | Santa Cruz |
| Phospho-JNK | Rabbit | 1:1000 | 4668 | Cell signaling |
| Total-JNK | Mouse | 1:1000 | sc-7345 | Santa Cruz |
| β-actin | Rabbit | 1:1000 | 4970 | Cell signaling |
|  | Mouse | 1:1000 | 004 | Abclonal |
| GAPDH | Rabbit | 1:1000 | sc-25778 | Santa Cruz |
|  | Mouse | 1:1000 | sc-32233 | Santa Cruz |
| Mouse IgG (IRDye 800CW) | Donkey | 1:1000 | 926-32212 | LI-COR |
| Rabbit IgG (IRDye 680RD) | Donkey | 1:1000 | 926-68073 | LI-COR |

Abbreviations: LXN, Latexin; PCNA, Proliferating cell nuclear antigen; ERK, Extracellular signal-regulated kinase; GAPDH, Glyceraldehyde-3-Phosphate Dehydrogenase; IgG, Immunoglobulin

### Proliferation assay

5-Bromo-2’-Deoxyuridine (BrdU) incorporation assay was performed in accordance with the manufacturer’s instruction (Millipore, #2752) ^18^. Briefly, 50×10^3^ cells in 200 μl of complete medium were seeded onto the 96-well plate and incubated overnight. After the starvation, cells were stimulated with PDGF-BB (315-18, PeproTech) or FBS in a dose-dependent manner with 25 μM of BrdU. Cells were washed, fixed, and incubated with a detector antibody for 1h. Then, cells were washed and incubated with a peroxidase-conjugated secondary antibody for 30 min. Finally, cells were incubated with 3, 3’, 5, 5’ - Tetramethylbenzidine (TMB) peroxidase substrate, and 450 nm wavelength was read by Synergy 2 microplate reader (Biotek).

### Transmigration assay

Transmigration assay was performed by Boyden’s chambers with polycarbonate membranes with 8 μm pores (Costar, #3422) as previously described ^12^. Briefly, the upper chamber was added with

0.5-1.0 × 10^6^ cells in 100 μl of serum-free conditional media. The lower chamber was added with or without PDGF-BB (20 ng/ml), FBS (10%), or MCP-1 (100 ng/ml) in 600 μl of serum-free conditional medium. After the stimulation, membranes were washed three times with cold PBS and fixed with methanol. Migrated cells on the lower side of the membrane were stained by crystal violet (Thermo Fisher Scientific, C581) and observed under a Nikon Eclipse 80i light microscope. The number of migrated cells through the membranes in three random fields was counted and averaged with ImageJ.

### Quantitative real-time polymerase chain reaction (qPCR)

Quantitative real-time polymerase chain reaction (qPCR) was performed as previously described^19^. Briefly, total RNAs were isolated and purified with TRIzol (Invitrogen, #15596026) in accordance with the manufacturer’s instructions. Complementary DNA (cDNA) was synthesized from 500 ng of total RNA using High Capacity cDNA Reverse Transcription Kit (Applied Biosystems, # 4368814). qRT-PCR analyses were conducted to quantitate the relative mRNA expression using SYBR Master Mix (Applied Biosystems, #4309155). Primer pairs used for qPCR are listed in Table 4. GAPDH was used as a housekeeping gene. After the reactions, the cycle threshold (Ct) data were determined using default threshold settings, and means Ct was determined from the triplicate PCRs. The PCR Ct values were normalized by subtracting the GAPDH Ct values, which provided the ΔCt value. A comparative ΔCt method was used to compare each condition with controls, and the values are expressed as 2^-ΔΔCt^.

**Table 4:**
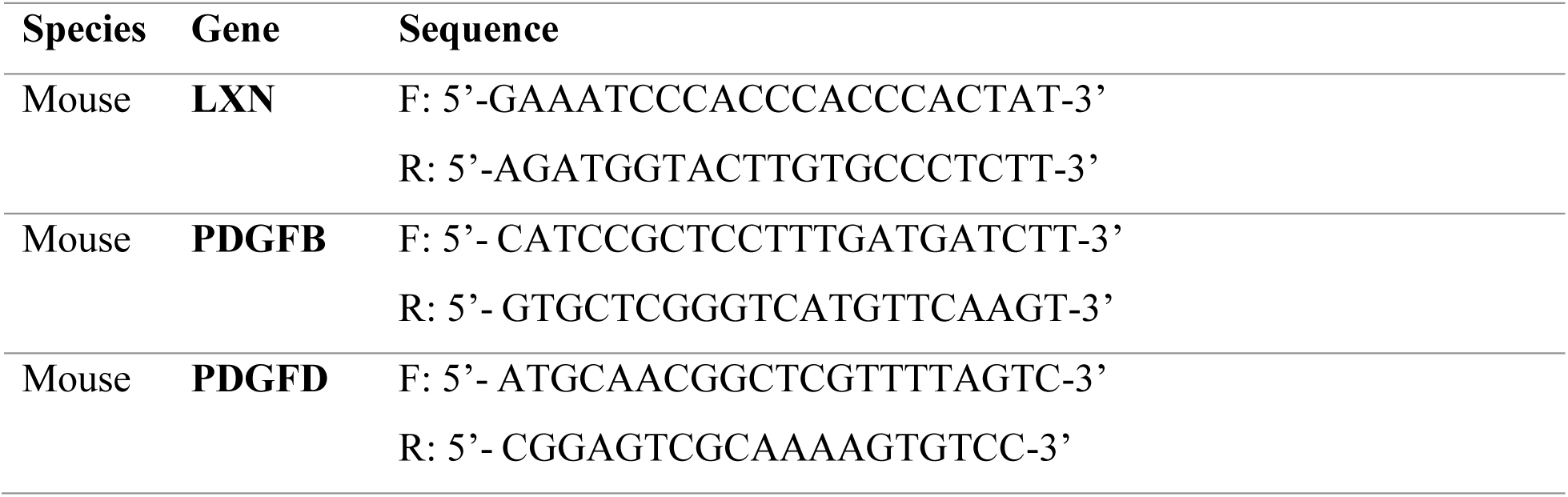
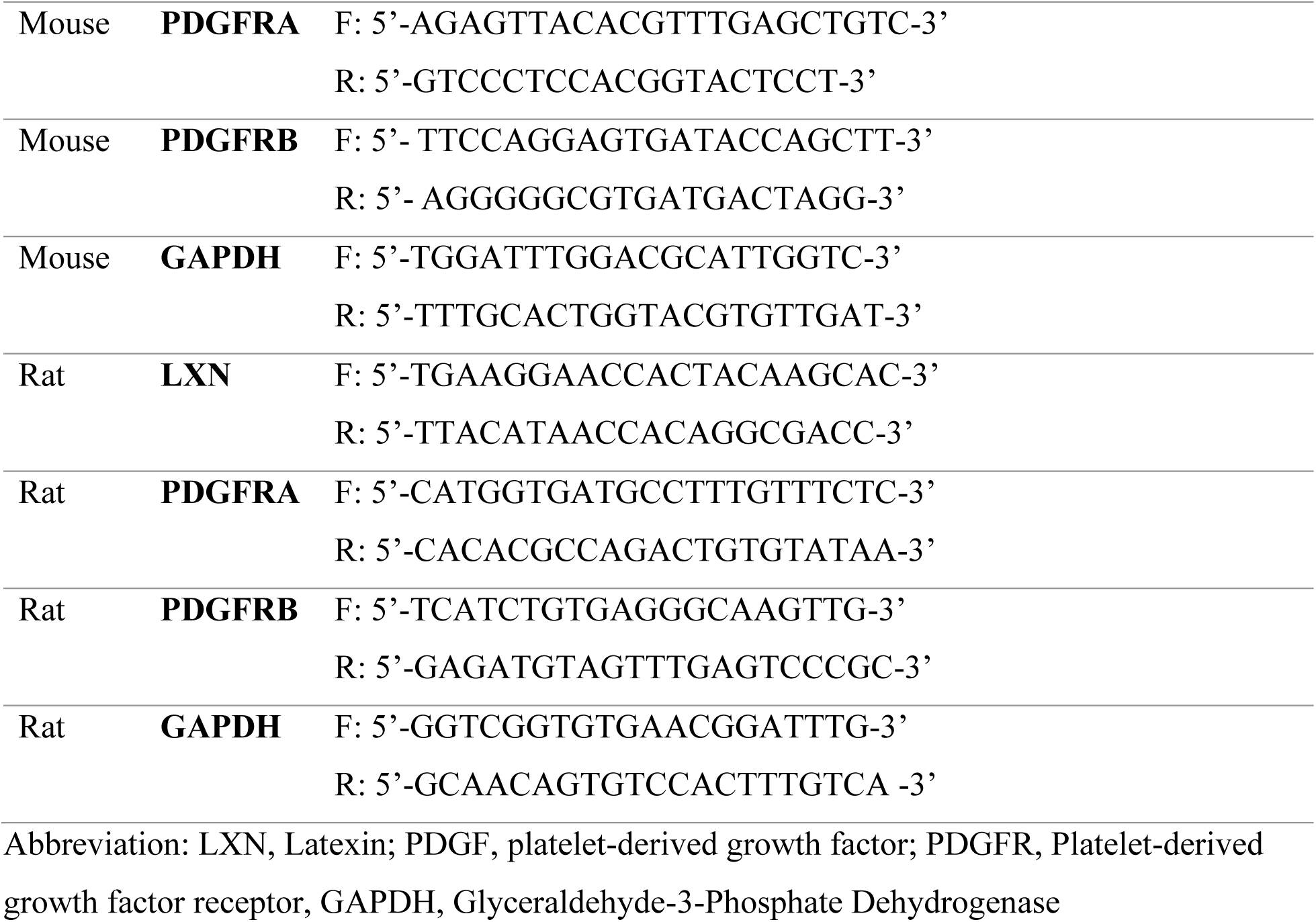
Primer pairs used for qPCR.

### Statistical analysis

All experiments were performed at least three times. All statistical analyses and graphs were performed and generated using GraphPad Prism software (version 5.0.1). Data were shown as means±SD. Statistical evaluations were performed by Student’s t-test for two groups or one-way analysis of variance (ANOVA) followed by Tukey’s test for multiple groups. Values of *P<0.05* were considered to be statistically significant, and the *P* value was described in the figures.

## Results

### LXN is significantly increased in carotid artery ligation-induced neointimal hyperplasia in mice

To understand the potential role of LXN in cardiovascular biology, we first investigated its expression in different tissues of mice. As shown in Fig. 1A, LXN is ubiquitously expressed in mouse tissues, as determined by western blot. The brain was one of the most highly LXN expressing tissue, which is consistent with previous notion that LXN is highly expressed in the lateral cortex ^4^. Furthermore, we found that LXN is also abundantly expressed in the aorta. Since LXN has been known as a potential pro-inflammatory protein ^20^, it indicates that LXN has a potential role in vascular pathology. To elucidate potential role of LXN in vascular inflammation, we examined whether LXN expression is regulated in a mouse model of neointimal hyperplasia after carotid artery ligation ^12^. As shown in Fig. 1B and 1C, our WB and IHC showed that LXN is substantially increased in the ligation injury group compared with the sham control group. To further investigate the pathological significance of LXN in neointimal hyperplasia, we next performed IHC co-staining with LXN and α-SMA, a contractile SMC marker, PCNA, a synthetic SMC marker, and CD68, a macrophage marker. As shown in Fig 1D, 1E, and 1F, α-SMA was decreased while PCNA and CD68 were increased in ligation group compared with sham group, indicating that increased SMC proliferation and macrophage infiltration are involved in this model, which is consistent with previous reports ^21–23^. Besides, we found that LXN was co-expressed with α-SMA and CD68, indicating that increased LXN is mainly localized in SMCs and macrophages. Taken together, these data suggest that LXN is involved in SMC proliferation and macrophage infiltration after vascular injury.

**Figure 1.**
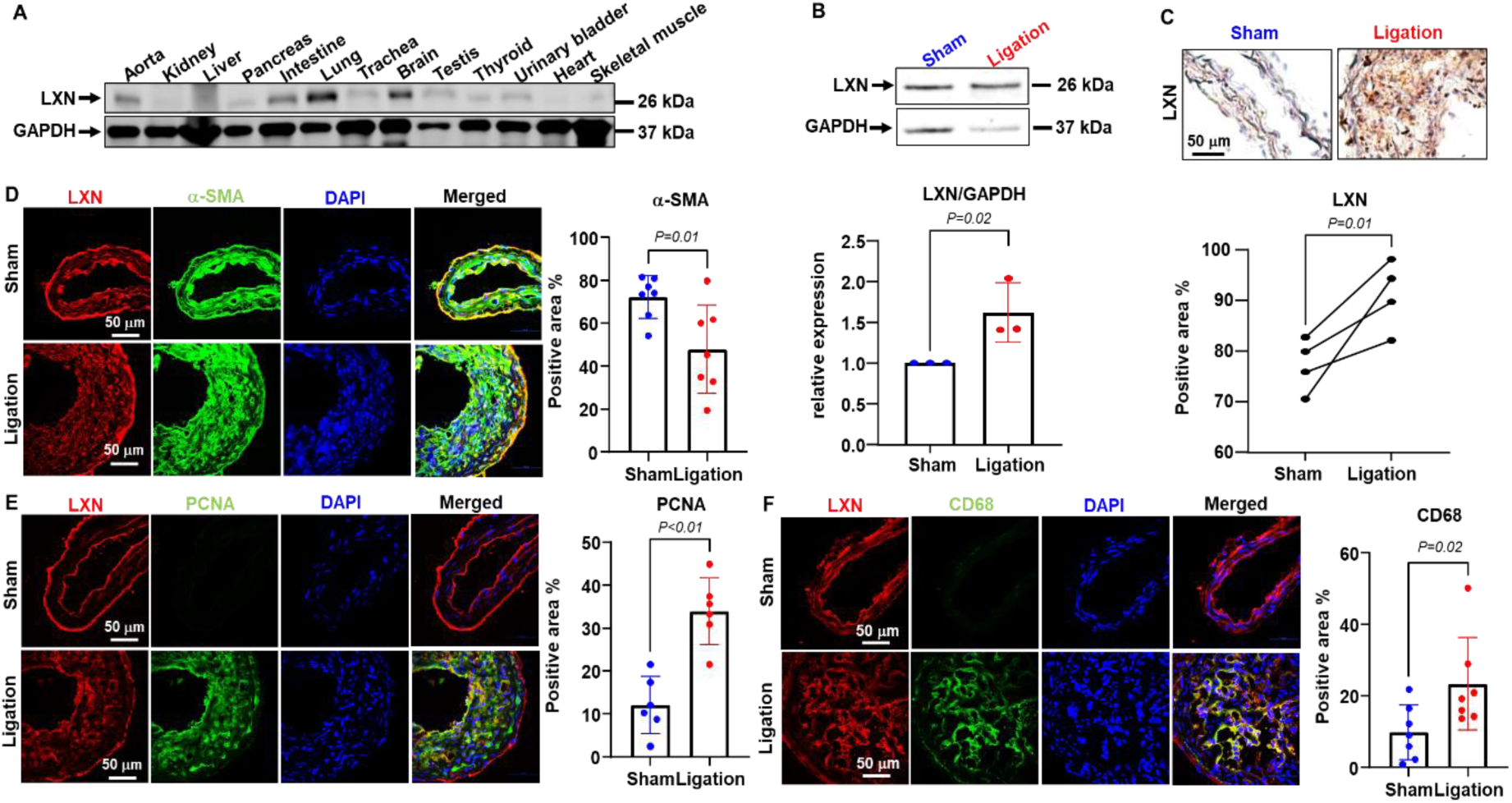
LXN is significantly increased in carotid artery ligation-induced neointimal formation in mice. A, Representative western blot (WB) image for LXN in different tissues of Wild type (WT) mice (n=3). GAPDH was used as internal control. B-F, WT mice were subjected to carotid artery ligation (left) or sham (right) surgery. 4-weeks after the surgery, carotid arteries were isolated and used for experiments. B, Representative WB and quantification for LXN (n=3). GAPDH was used as internal control. C, Representative immune-histochemistry (IHC) and quantification for LXN (n=4). Scale bar=100 μm. D-F, Representative IHC co-staining and for LXN (red) and α-SMA (green) (D, n=7) or PCNA (green) (E, n=6), or CD68 (F) (n=7). DAPI (blue) was used for nuclei staining. Positive area % was quantified by Image J software. Scale bar=50 μm.

### Global LXN deficiency suppresses carotid artery ligation-induced neointimal hyperplasia in mice

To further investigate the role of LXN in vascular remodeling, we next examined whether LXN deficiency prevents carotid artery ligation-induced neointimal hyperplasia. We performed the sham or ligation surgery on LXN global KO mice (LXN-gKO) or their wild-type littermates (LXN-gWT). As shown in Fig. 2A, our histology demonstrated that LXN-gKO strikingly prevents neointimal hyperplasia compared with LXN-gWT at 4 weeks after injury. Similar results were obtained after performing serial sectioning of the carotid arteries harvested from LXN-gWT and - gKO mice after 4-week injury (Supplemental fig IA-ID). Further, as shown in Fig. 2B-2D, our IHC suggested that LXN-gKO increases α-SMA expression and decreases PCNA and CD68 expressions in the neointimal lesions, indicating that LXN may prevent neointimal hyperplasia through regulating SMC proliferation and macrophage infiltration.

**Figure 2.**
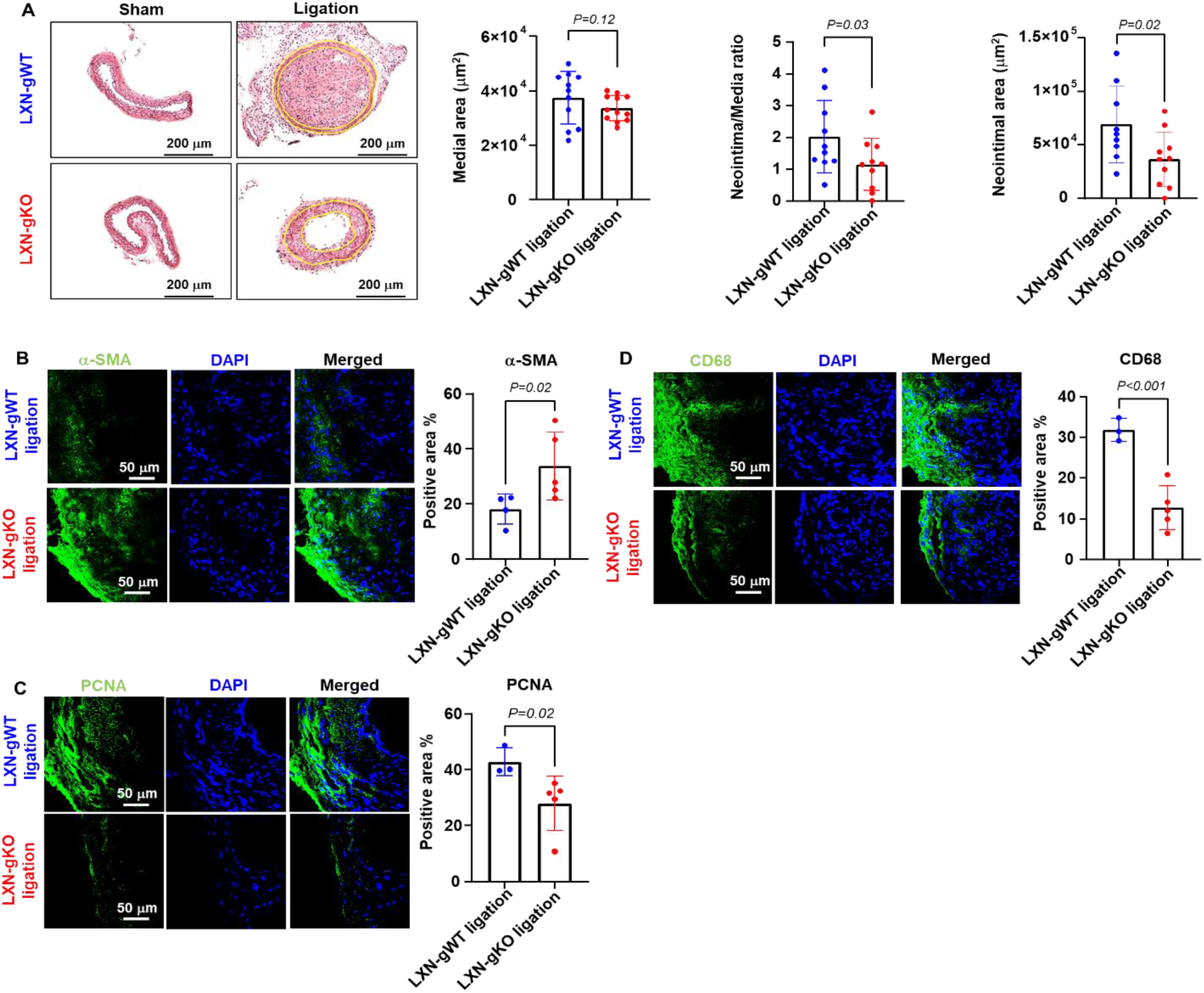
Global LXN deficiency suppresses carotid artery ligation-induced neointimal hyperplasia in mice. Global LXN knockout (LXN-gKO) or its control (LXN-gWT) mice were subjected to carotid artery ligation (left) or sham (right) surgery. 4-weeks after the surgery, carotid arteries were isolated and used for experiments. A, Representative HE staining images are shown and Medial area (n=11-12), Neointimal area (n=10), and Neointima/Media ratio (n=9-10) was measured by Image J software. Scale bar=200 μm. B-D, Representative IHC staining for α-SMA (green) (B, n=4-5), PCNA (green) (C, n=3-5), or CD68 (green) (D, n=3-5). DAPI (blue) was used for nuclei staining. Positive area % was quantified by Image J software. Scale bar=50 μm.

### SMC-specific LXN deficiency suppresses carotid artery ligation-induced neointimal hyperplasia in mice

LXN was previously identified as a laminar shear stress responsive gene in ECs (Ref). Since endothelial cell (EC) activation induced vascular inflammation is considered as a primary event of neointimal hyperplasia^24^, we then investigated whether EC-specific LXN deficiency can prevent carotid artery ligation-induced neointimal hyperplasia. EC-specific LXN-KO (LXN-eKO) and its control mice (LXN-eWT) using VE-cadherin Cre/ERT2 recombinase with a floxed LXN allele were generated (Supplemental fig IIA). As shown in Supplemental fig IIB, LXN expression is significantly reduced in CD31-positive cells isolated from LXN-eKO lungs as compared with LXN-eWT after tamoxifen injection (75 mg/kg, intraperitoneal, 5 consecutive days), whereas LXN expression in CD31-negative lung cells remains the same, as determined by western blot. Furthermore, we found that LXN-eKO did not prevent carotid artery ligation-induced neointimal hyperplasia (Supplemental fig IIC). These data indicate that endothelial LXN has no or minimal effects on neointimal formation.

To investigate whether SMC LXN is involved in neointimal formation, we generated SMC-specific LXN-KO mice (LXN-sKO) using SM22 Cre recombinase with a floxed LXN allele. LXN-smcKO were identified by genotyping using PCR (Fig 3A-C). LXN knockout efficiency was confirmed by WB and qPCR. As shown in Fig. 3D-3G, LXN was efficiently knockdown in the mouse aorta as well as in primarily cultured SMCs. LXN-sWT and -sKO mice were then subjected to carotid artery ligation for 4-weeks. As shown in Fig. 3H, we found that LXN-sKO substantially prevents carotid artery ligation-induced neointimal hyperplasia compared with LXN-sWT as determined by histological staining. Moreover, as shown in Fig 3I and 3J, LXN-sKO led to an increased expression of α-SMA, a contractile SMC marker, and decreased expression of PCNA, a synthetic SMC marker, indicating that LXN-sKO prevents SMC proliferation. Taken together, these data suggest that LXN in SMCs plays a role in neointimal formation after ligation injury.

**Figure 3.**
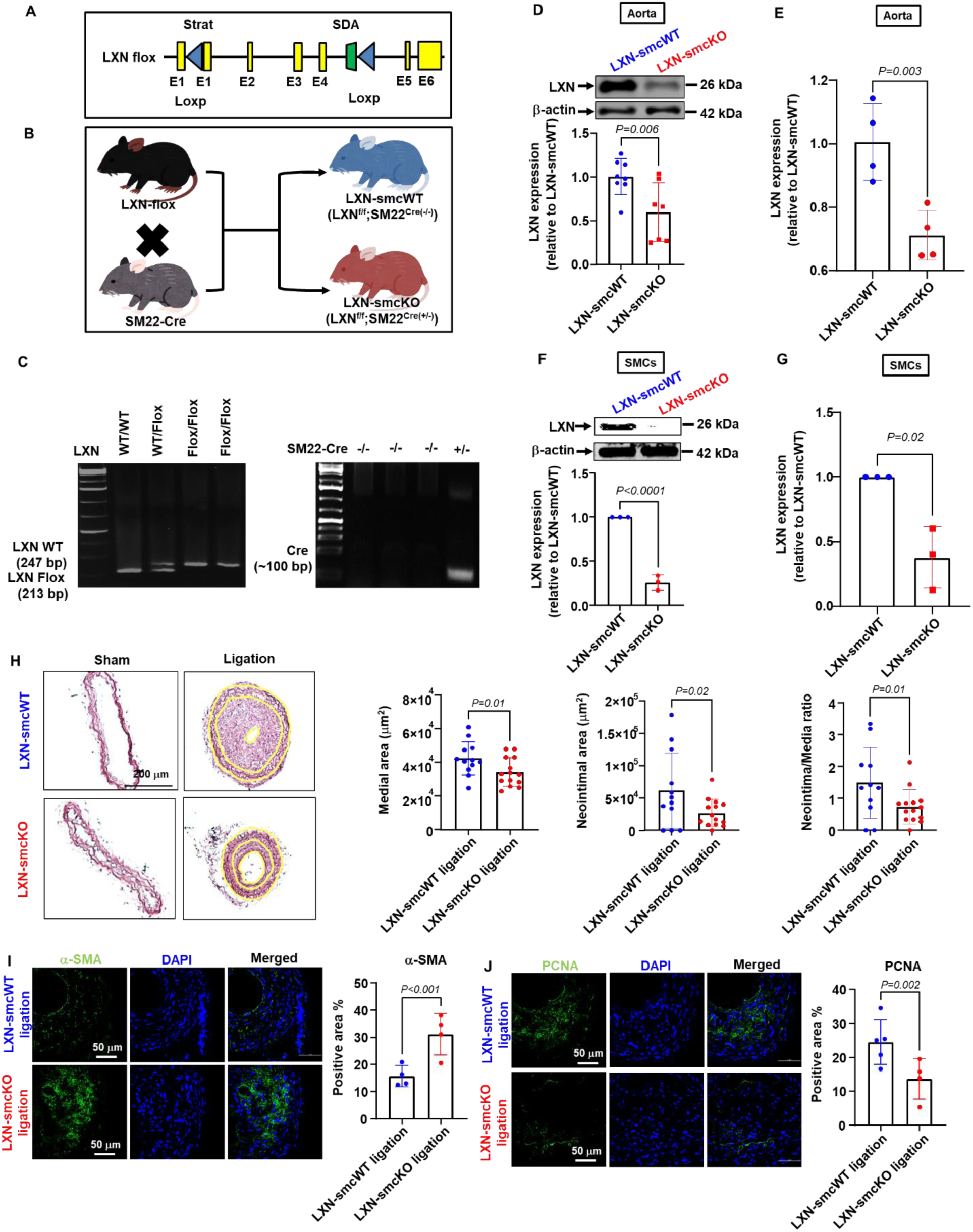
SMC-specific LXN deficiency suppresses carotid artery ligation-induced neointimal formation in mice. A, Illustration of genotyping strategy. B, Illustration of breeding outcome for SMC-specific LXN KO mice. C, Representative genotyping PCR images for LXN-Flox and SM22-Cre. D and E, Representative WB and quantification (D, n=7-8) or quantitative polymerase chain reaction (qPCR; E, n=4) for LXN with thoracic aortas isolated from LXN-sWT (LXN^flox/flox^; SM22^cre(–/–)^) and-sKO (LXN^flox/flox^; SM22^cre(+/–)^) mice. β-actin was used as internal control for WB. F and G, Representative WB and quantification (F, n=3) or qPCR (G, n=3) for LXN with primary SMCs isolated from LXN-sWT and-sKO mice. β-actin was used as internal control for WB. H-J, LXN-sWT and -sKO mice were subjected to carotid artery ligation (left) or sham (right) surgery. 4-weeks after the surgery, carotid arteries were isolated and used for experiments. H, Representative HE staining images. Scale bar=200 μm. Medial area, Neointimal area, and Neointima/Media ratio were measured by Image J software (n=12-14). I-J, Representative IHC and quantification for α-SMA (I, green, n=4) and PCNA (J, green, n=4-5). DAPI (blue) is used for nuclei staining. Scale bar=50 μm. Positive area % was quantified by Image J software.

### LXN deficiency suppresses FBS-induced mouse SMC proliferation and migration

To further identify the potential mechanisms of regulating SMC biology by LXN, we next performed in vitro assays with mouse primary SMCs. First, we examined whether LXN expression is changed in response to FBS. As shown in Fig. 4A, our WB demonstrated that LXN is significantly increased in response to FBS (48 h) treatment in a dose-dependent manner. PCNA, which is a marker for cell proliferation, is induced in response to FBS treatment. Then, we next investigated whether LXN deficiency prevents SMC proliferation and/or migration, which are critically involved in neointimal hyperplasia ^1^. As shown in Fig. 4B, our aortic explant assay demonstrated that LXN deficiency markedly prevented SMC outgrowth. Since the aortic explant assay includes both SMC proliferation and migration, we next performed cell proliferation and migration assays. As shown in Fig 4C and 4D, LXN deficiency prevents FBS-induced SMC proliferation, as determined by cell counting and BrdU incorporation assays, and FBS-induced SMC migration (Fig. 4E), as determined by transwell migration assays. Taken together, our results demonstrated that LXN is increased in proliferative SMCs, and its deficiency prevents FBS-induced SMC proliferation and migration.

**Figure 4.**
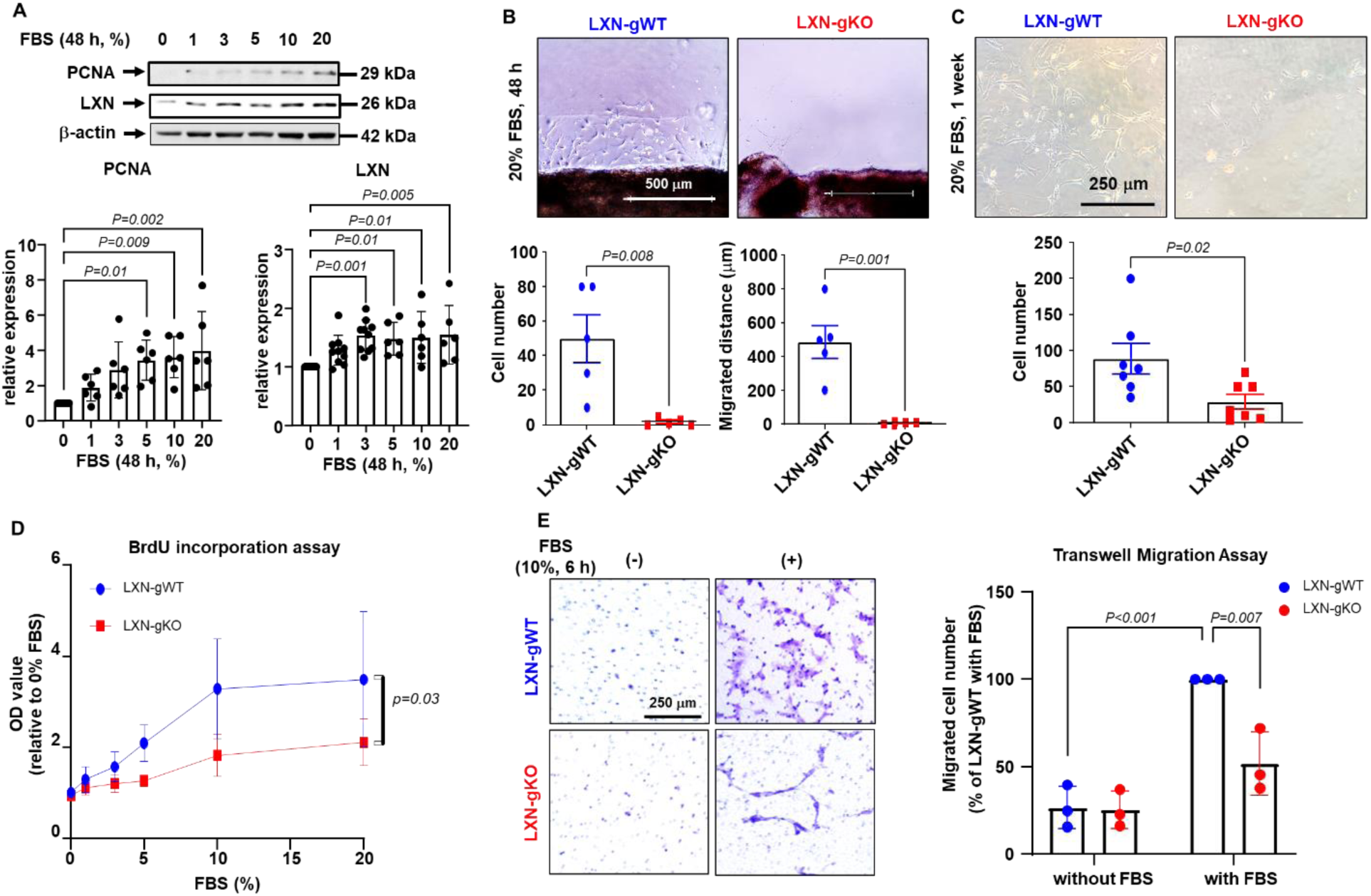
LXN deficiency suppresses proliferation and migration in mouse SMCs. A, Representative WB images and quantification for PCNA and LXN. Primary SMCs isolated from WT mice were treated with FBS for 48 hours in a dose-dependent manner (0-20 %) (n=6). β-actin was used as internal control. B, Representative images and quantification of explant aortic tissue culturing of LXN-gWT and -gKO mice for 48 hours with 20% fetal bovine serum (FBS) (n=5). Scale bar=500 μm. C, Representative images and quantification of primary SMC proliferation isolated from LXN-gWT and -gKO mice. Mouse primary SMCs were cultured for 1 week with 20% FBS (n=7). Scale bar=250 μm. D, BrdU incorporation assay with primary SMCs isolated from LXN-gWT and -gKO mice. Primary SMCs were treated with FBS for 24 h in a dose-dependent manner (0-20%) (n=4-6). E, Representative images and quantification of Transwell migration assay with primary SMCs isolated from LXN-gWT and -gKO mice. Primary SMCs were treated with or without 10% FBS for 6 h. Scale bar=250 μm. Migrated cell number was counted by image J software (n=3).

### LXN deficiency suppresses PDGF receptor expressions in mouse SMCs

Furthermore, to identify the molecular targets in regulating SMC proliferation by LXN, we performed bulk RNA-sequencing in SMCs isolated from LXN-gKO and their WT littermates. We found that several proliferative related genes or pathways, such as KLFs, PDGF/PDGFRs, collagen, and TGF-beta and IGF, were differentially expression in WT and LXN-deficient SMCs. We further confirmed the expression of these genes by qPCR. We found that LXN deficiency substantially attenuated the expression of PDGFRA and PDGFRB in SMCs as compared with SMCs derived from WT mice as shown in Fig 5A and 5B, whereas other genes, such as CCND1, ITGAs, KLFs, and TGF-1β, were not significantly affected by LXN knockout in SMCs (Supplemental fig III). Taken together, our data strongly suggest that LXN deficiency may attenuate the expression PDGFR in mouse primary SMCs, which may contribute to the decreased proliferation and migration observed in LXN deficient vascular SMCs.

**Figure 5.**
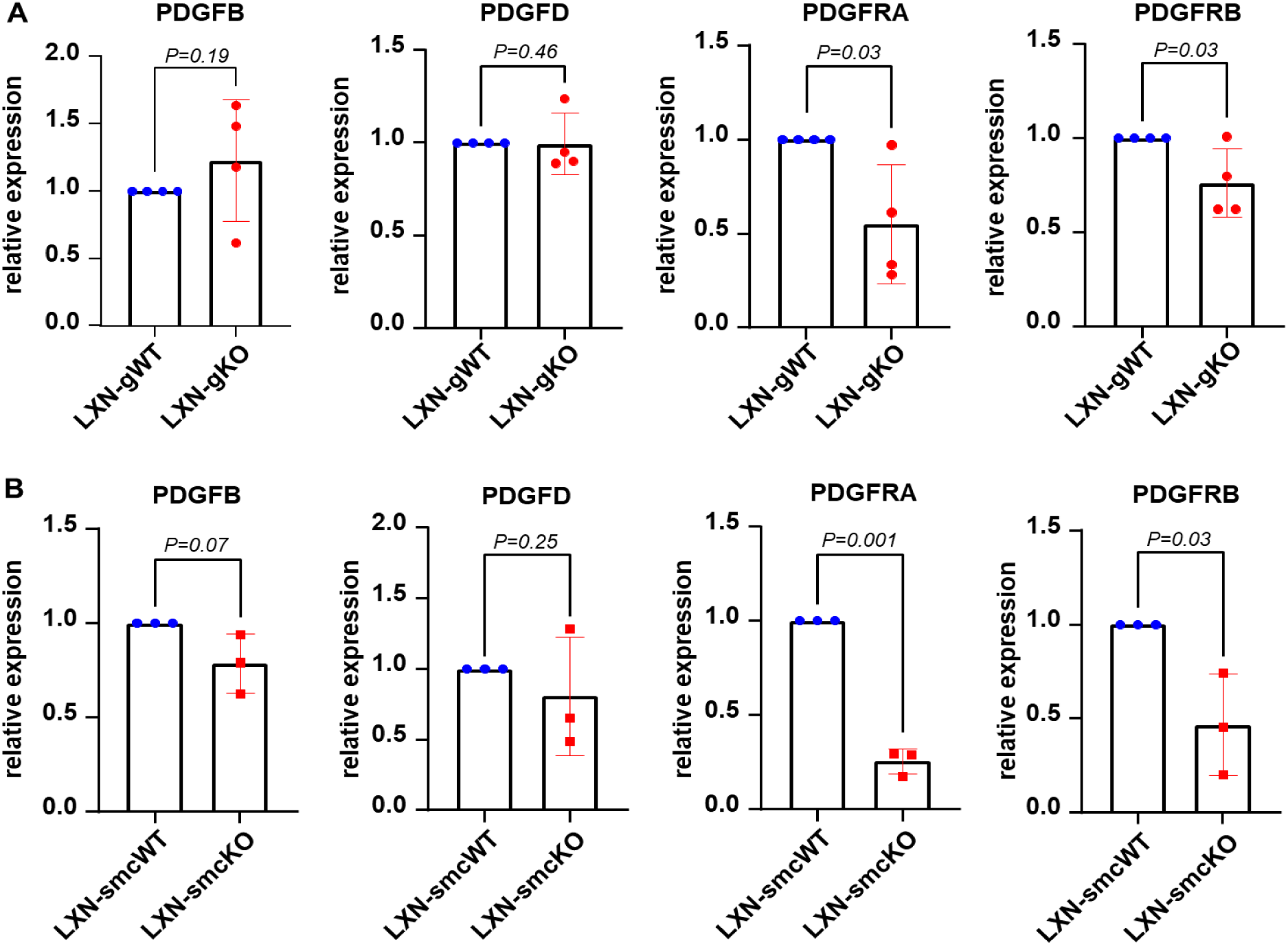
LXN deficiency suppresses PDGF receptor expressions in mouse SMCs. A-B, qPCR for PDGFB, PDGFD, PDGFRA, and PDGFRB with primary SMCs isolated from LXN-gWT and -gKO (A, n=4) or LXN-sWT and -sKO mice (B, n=3).

### LXN deficiency suppresses PDGF-BB-induced rat SMC proliferation and migration

To further investigate the role of PDGFR in regulating SMC proliferation by LXN, we tested whether LXN deficiency prevents PDGF-BB-dependent SMC proliferation and migration and its signaling pathways. To this end, rat aortic SMCs were transfected with small hairpin RNA (shRNA) for rat LXN or Non-Target control. As shown in Fig. 6A and 6B, LXN-KD prevented PDGF-BB-induced PCNA expression and BrdU incorporation, indicating that LXN-KD prevents PDGF-BB-induced SMC proliferation. Also, as shown in Fig. 6C, our transwell migration assay suggested that LXN-KD prevents PDGF-BB- and FBS-induced SMC migration. Mechanistically, as shown in Fig. 6D, our WB demonstrated that LXN-KD remarkably attenuates PDGF-BB-induced phosphorylation of mitogen-activated protein kinase (MAPK) pathways, such as ERK, p38, and JNK, whereas phosphorylation of NF-κB and Akt were not affected. Likewise, LXN-KD significantly attenuated PDGFRA expression (Fig. 6E), which is notably consistent with our data obtained in mouse primary SMCs (Fig. 5A and 5B). Taken together, our results suggest that LXN deficiency prevents PDGF-BB-dependent SMC proliferation and MAPK phosphorylation mainly through attenuating expression of PDGF receptor expression (Fig. 6F).

**Figure 6.**
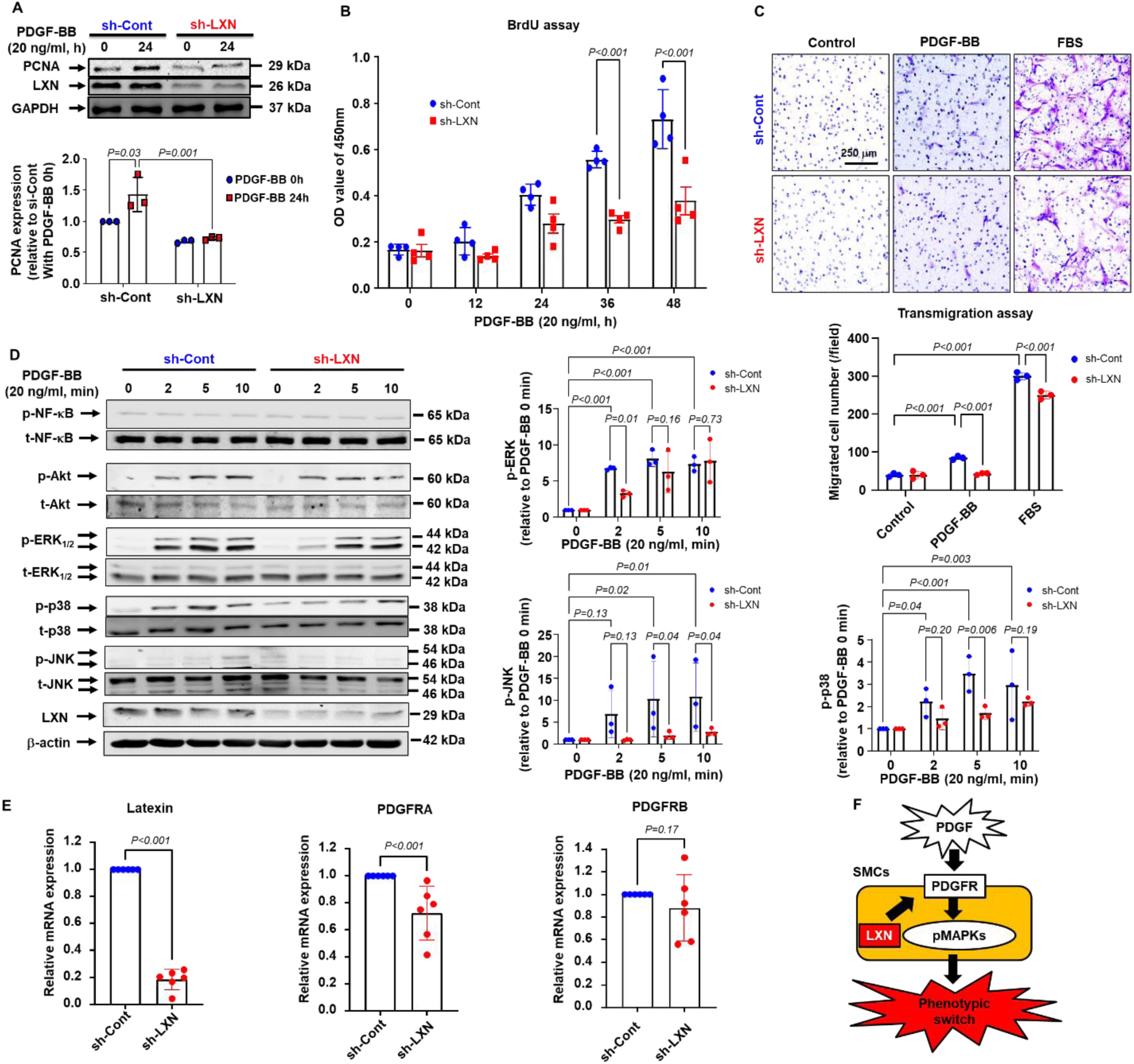
LXN deficiency suppresses PDGF-BB-induced proliferation and migration in rat SMCs. Rat aortic SMCs were pretreated with sh-Control or sh-LXN and then stimulated with or without Platelet-derived growth factor (PDGF)-BB. A, Representative WB and quantification for PCNA with rat SMCs simulated with or without PDGF-BB (20 ng/ml, 24 h) in the presence of sh-Control (sh-Cont) or sh-LXN (n=3). GAPDH was used as internal control. LXN was used to confirm knockdown efficiency. B, BrdU incorporation assay with rat SMCs stimulated with PDGF-BB (20 ng/ml) in a time-dependent manner (0-48 h) in the presence of sh-Control or sh-LXN (n=4). C, Representative transmigration images and quantification of migrated cell numbers. Rat SMCs pretreated with sh-control or sh-LXN were stimulated with the vehicle, PDGF-BB (20 ng/ml), or FBS (10%) for 24 h. Scale bar=250 μm. Migrated cell number was counted by image J software (n=3). D, Representative WB and quantification for phospho-Akt (p-Akt), and phospho-NF-κB (p-NF-κB), phospho-ERK 1/2 (p-ERK 1/2), phospho-p38 (p-p38), and phospho-pJNK (p-pJNK) (n=3). Total-Akt (t-Akt), total-NF-κB (t-NF-κB), total-ERK 1/2 (t-ERK 1/2), total-p38 (t-p38), total-JNK (t-JNK), and β-actin was used as an internal control. E, qPCR for LXN, PDGFRA, and PDGFRB (n=6). E, Schematic figure of LXN function in SMCs.

### Myeloid-specific LXN deficiency suppresses carotid artery ligation-induced neointimal hyperplasia in mice

It has been reported that LXN plays a pro-inflammatory role in macrophages ^20^. Further, as shown in Fig. 2A and 2D, LXN-gKO significantly prevents neointimal formation and macrophage infiltration. Therefore, we attempted to investigate whether LXN in macrophages plays a role in in neointimal formation. To this end, we generated myeloid-specific LXN KO mice (LXN-mKO) and their control mice (LXN-mWT) by crossing Lyz2 Cre mice with LXN floxed mice (Fig. 7A and 7B). As shown in Fig. 7C, our WB demonstrated that LXN expression is specifically knockout in peritoneal macrophages (PMs) obtained from LXN-mKO compared with LXN-mWT, whereas LXN expression in the aorta was not altered. Importantly, as shown in Fig. 7D, LXN-mKO significantly prevented carotid artery ligation-induced neointimal hyperplasia compared with WT littermates after carotid artery ligation for 4 weeks. Likewise, CD68-positive area in neointimal lesions was markedly attenuated in LXN-mKO mice compared with LXN-mWT mice. Taken together, these data suggest that LXN in macrophages contributes to neointimal formation after injury.

**Figure 7.**
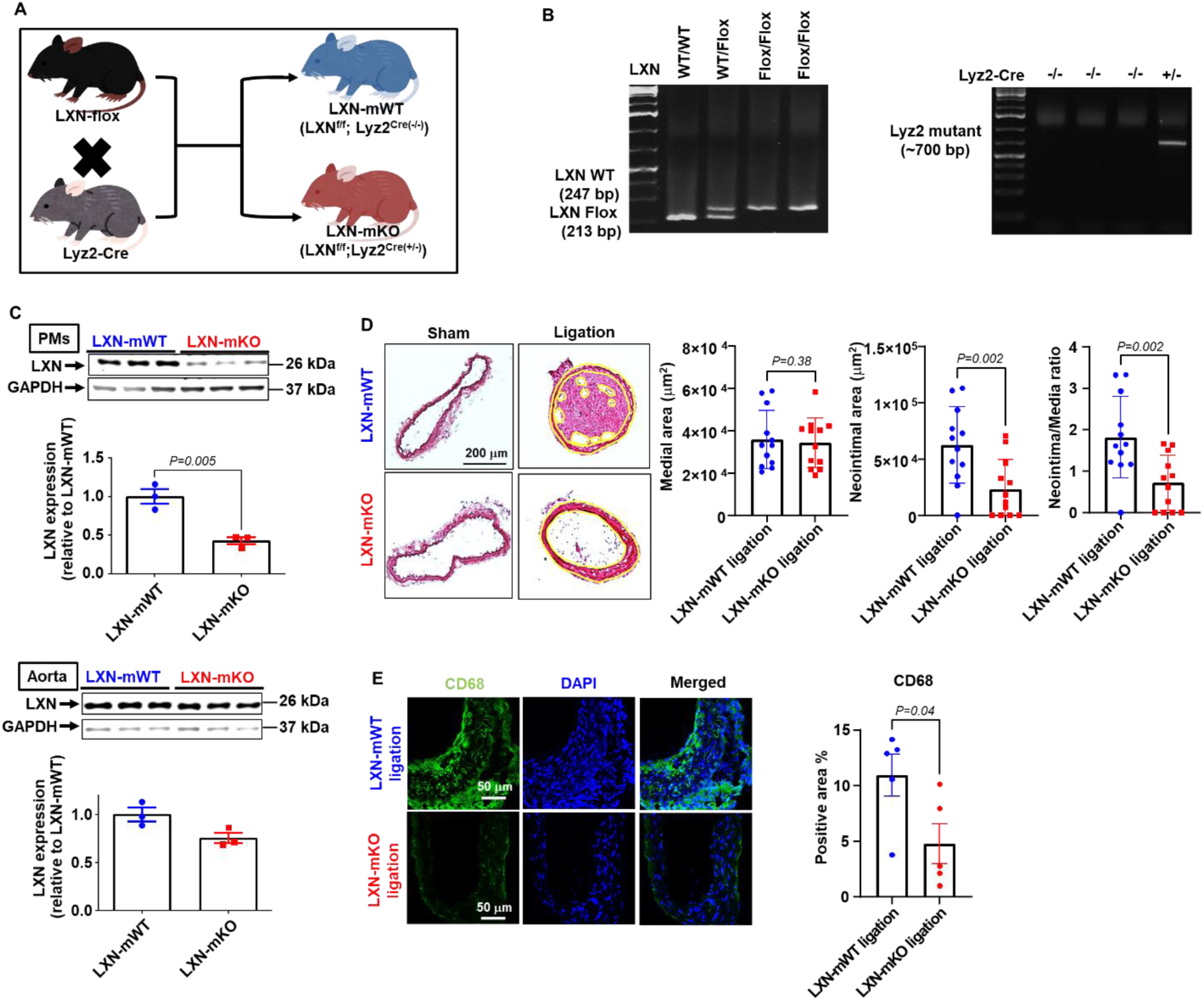
Myeloid-specific LXN deficiency suppresses carotid artery ligation-induced neointimal hyperplasia in mice. Myeloid-specific LXN-KO (LXN^flox/flox^, Lyz-^Cre^ (+/–); LXN-mKO) and its control mice (LXN^flox/flox^, Lyz-^Cre (–/–)^; LXN-mWT) were used. A, Illustration of breeding outcome for LXN-mKO mice. B, Representative genotyping PCR image for LXN-Flox and Lyz-cre. C, Representative WB and quantification for LXN with peritoneal macrophages (PMs) or thoracic aortas isolated from LXN-mWT or -mKO (n=3). GAPDH was used as internal control. LXN-mWT and -mKO mice were subjected to carotid artery ligation (left) or sham (right) surgery. 4-weeks after the surgery, carotid arteries were isolated and used for experiments. D, Representative HE staining images. Scale bar=200 μm. Medial area, Neointimal area, and Neointima/Media ratio were measured by Image J software (n=12). E, Representative IHC and quantification for CD68 (green). DAPI (blue) is used for nuclei staining. Scale bar=50 μm. Positive area % was quantified by Image J software (n=5).

### LXN deficiency prevents MCP-1-induced macrophage migration through inhibition of ERK phosphorylation in vitro

Since MCP-1-induced macrophage migration is a critical event for neointimal hyperplasia ^25^, we next examined whether LXN deficiency prevents MCP-1-induced macrophage migration. As shown in Fig. 8A, our transwell migration assays demonstrated that LXN-gKO significantly prevented MCP-1-induced PM migration. Mechanistically, as shown in Fig. 8B, MCP-1-induced ERK phosphorylation is dramatically attenuated in PMs obtained from LXN-gKO compared with LXN-gWT. Similarly, LXN-mKO significantly prevented MCP-1-induced migration and ERK phosphorylation in PMs (Fig. 8C and 8D). To verify the role of ERK phosphorylation in PM migration, we next examined whether a pharmacological ERK inhibitor U0126 prevents MCP-1-induced PM migration. As shown in Supplemental fig V, U0126 efficiently inhibits MCP-1-induced ERK phosphorylation in PMs. Importantly, as shown in Fig. 8E, U0126 completely prevented MCP-1-induced PM migration, indicating that ERK phosphorylation is essentially involved in MCP-1-induced PM migration. These data indicate that LXN deficiency prevents MCP-1-induced ERK phosphorylation, which is critically involved in macrophage migrationand neointimal hyperplasia (Fig 8F).

**Figure 8.**
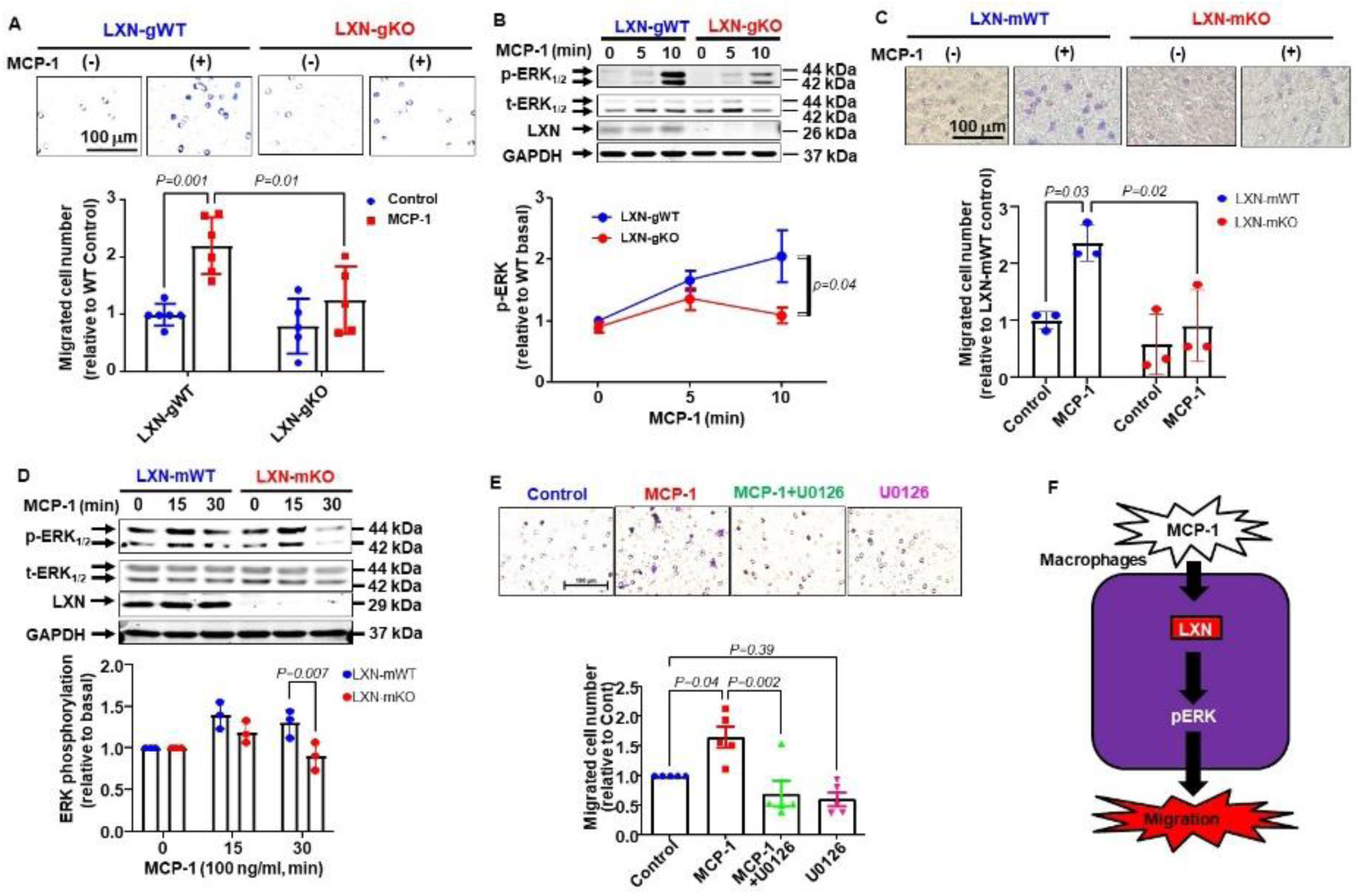
LXN deficiency prevents MCP-1-induced macrophage migration through inhibition of ERK phosphorylation in vitro. A, Representative transmigration images and quantification of migrated cell numbers. PMs isolated from LXN-gWT or -gKO were stimulated with or without monocyte chemoattractant protein-1 (MCP-1; 100 ng/ml, 6 h). Scale bar=100 μm. Migrated cells were counted by image J software (n=5-6). B, Representative WB and quantification for phospho-ERK 1/2 (p-ERK 1/2). PMs isolated from LXN-gWT or -gKO were stimulated with MCP-1 (100 ng/ml) in a time-dependent manner (0-10 min) (n=3). Total-ERK 1/2 (t-ERK 1/2) and GAPDH was used as internal control. LXN was used to confirm knockout efficiency. C, Representative transmigration images and quantification of migrated cell numbers. PMs isolated from LXN-mWT or -mKO were stimulated with or without MCP-1 (100 ng/ml, 6 h) (n=3). Scale bar=100 μm. D, Representative WB and quantification for p-ERK 1/2. PMs isolated from LXN-mWT or -mKO were simulated with MCP-1 (100 ng/ml) in a time-dependent manner (0-30 min) (n=3). T-ERK 1/2 and GAPDH was used as internal control. LXN was used to confirm knockout efficiency. E, Representative transmigration images and quantification of migrated cell numbers. PMs isolated from WT mice were stimulated with or without MCP-1 (100 ng/ml, 6 h) in the absence or presence of U0126 (10 μM, 24 h), an ERK inhibitor (n=5). Scale bar=100 μm. Migrated cells were counted by Image J software. F, Schematic figure of LXN function in Macrophages.

## Discussion

Although it has been known that vascular inflammation, SMC phenotypic switch, and macrophage infiltration are involved in neointimal hyperplasia and restenosis ^1^, the detailed underlying mechanisms are not fully understood. Since LXN is highly expressed in vascular wall and its expression is responsive to laminar flow, we investigated the role of LXN roles in vascular remodeling after injury. In the present study, we identified LXN as an essential regulator in the development of neointimal hyperplasia after vascular injury. We found that LXN is highly expressed in neointimal lesions and that both SMC-and macrophage-deficient mice markedly attenuated neointimal formation in a mouse model of carotid artery ligation. Mechanistically, we found that LXN regulates PDGF-BB-induced SMC proliferation and MCP-1-induced macrophage migration through attenuating PDGFRA expression in SMCs and MCP-1-induced ERK phosphorylation in macrophages.

LXN has been shown to ubiquitously expressed in different cells and organs, such as neural cells, immune-cells, cancer tissues, and hematopoietic stem cells ^5, 9^ ^6^ ^8^ ^9^ ^20^. Although LXN has been known as a carboxypeptidase A inhibitor ^20^, previous studies have been mainly focused on its non-canonical roles in inflammation and stem cell proliferation. For instance, LXN expression is induced in LPS-stimulated macrophages (ref). Indeed, recent studies have implicated LXN in the development of several inflammatory diseases, such as acute pancreatitis and alcoholic hepatitis. Additionally, Ying Liang et al. have shown that LXN is a negative stem cell regulatory gene through regulating stem cell proliferation and apoptosis ^6–8^. Although LXN is highly expressed in blood vessels and its expression is dramatically downregulated in ECs in response to laminar shear stress, the role of LXN in regulating vascular hemostasis has not been explored thus far. As far as we know, this is the first study to systemically investigate the cell-specific roles of LXN in vascular remodeling.

EC inflammation is considered as a primary event of neointimal hyperplasia ^24^, we first investigated whether EC-specific LXN deficiency prevents neointimal hyperplasia. LXN was abundantly expressed in ECs; however, its deficiency in ECs did not prevent neointimal hyperplasia (Supplemental fig. II). On the other hand, accumulating evidence has revealed the roles of the vascular SMCs in the development of cardiovascular diseases, such as restenosis and atherosclerosis. In damaged blood vessels, contractile SMCs are dedifferentiated into synthetic phenotypes (proliferative and migratory) in response to pro-inflammatory cytokines and growth factors, called SMC phenotypic switch ^1^ ^2^ ^21–23^. Synthetic SMC is defined as a combination of lower expression of α-SMA and higher expression of PCNA ^2^, which pattern was observed in the ligation group in the present study (Fig 1D and 1E), indicating that SMC phenotypic switch is definitely involved in our neointimal hyperplasia model. Additionally, our IHC identified that LXN is co-expressed with α-SMA and PCNA (Fig. 1D and 1E). From these data, we next hypothesized that SMC-specific LXN deficiency attenuates neointimal hyperplasia. In fact, SMC-specific LXN deficiency prevented carotid artery ligation-induced neointimal formation (Fig. 3H) as well as SMC phenotypic switch (Fig. 3I and 3J). In mouse SMCs, LXN was induced in response to FBS treatment (Fig. 4A) and its deficiency prevented FBS-induced proliferation (Fig. 4C and 4D) and migration (Fig. 4E). Importantly, LXN deficiency suppressed PDGF receptor expressions (Fig. 5A and 5B). Moreover, LXN deficiency prevented PDGF-BB-induced proliferation (Fig. 6A and 6B) and migration (Fig. 6C) through inhibition of MAPK phosphorylation (Fig. 6D) attenuating PDGF receptor expressions (Fig. 6E) in rat SMCs. These data indicates that LXN regulates PDGF receptor expressions in SMCs, which is critically involved in carotid artery ligation-induced neointimal hyperplasia and SMC phenotypic switch; however it remains unknown how LXN regulates PDGF receptor expressions. Since LXN is previously considered as a cytosolic protein with limited access to the nuclear ^27^, LXN may not exert its function here as a nuclear factor. On the other hand, PDGF receptor expressions are regulated by growth factors, such as transforming growth factor (TGF) and PDGF itself ^28^. Since our WB suggested that LXN plays a role in PDGF-induced MAPK phosphorylation (Fig. 6D), this might be involved in a negative feedback of PDGF receptor expressions. Moreover, LXN is recently considered as a secretory protein having an inflammatory role ^27^. Based on this report, extracellular LXN may regulate PDGF receptor expressions through autocrine, paracrine, and endocrine signaling as a cytokine. Further investigation is needed to determine how LXN regulates PDGF receptor expressions.

Consistent with previous notion that LXN may play an inflammatory role in macrophages ^20^, we found that LXN is co-expressed not only in SMCs but also in CD68-positive macrophages in carotid artery ligation-induced neointimal hyperplasia (Fig. 1E) and that global LXN deficiency prevented neointimal hyperplasia (Fig. 2A) and macrophage infiltration (Fig. 2D). Since macrophage infiltration contributes to various cardiovascular disease ^21–23^, we next investigated whether macrophage-specific LXN deficiency prevents neointimal hyperplasia in mice. As hypothesized, our histology demonstrated that myeloid-specific LXN deficiency strikingly prevents carotid artery ligation-induced neointimal hyperplasia (Fig. 7D) as well as macrophage infiltration (Fig. 7E). In macrophages, LXN deficiency prevented MCP-1-induced migration through inhibition of ERK phosphorylation (Fig 8). The molecular mechanisms underlying regulating ERK phosphorylation remains elusive. Recently, Heinrich et al. have reported that LXN is highly correlated to the M1 macrophage phenotype ^29^. M1 macrophages are considered to be pro-inflammatory phenotype expressing higher levels of MCP-1 receptor, CCR2, whereas M2 are more anti-inflammatory ^30, 31^. Based on these, macrophage in LXN deficient mice could be more anti-inflammatory M2 phenotypes, which express less CCR2 levels with decreased phosphorylation of ERK. Further studies are needed to investigate how LXN regulates macrophage phenotype and infiltration.

Although we have characterized a role of LXN in neointimal hyperplasia, the distribution of LXN expression in vasculature remains unclear. LXN is highly induced in neointimal hyperplasia in vivo, whether this increase occurs inside the cells or in extracellular matrix remains to be determined. In our study, we found that increased LXN expression in SMCs seems to be an agonist dependent. For instance, LXN was slightly but statistically significantly induced in response to FBS treatment in mouse SMCs (Fig 4A) in vitro, whereas PDGF treatment did not change LXN levels in rat SMCs (Fig 6C). Similarly, although a previous report has shown that LXN is induced in response to LPS and CSF-1 treatments in macrophages ^20^ , in the present study, LXN expression was not altered in response to MCP-1 treatment in macrophages (data not shown). Recent studies suggest that LXN is considered as a secretory protein ^27^, thus LXN might be chronically secreted from vascular cells and immune cells under pathological conditions, which could be involved in increased LXN levels in the lesions in vivo. Further studies are needed to investigate LXN expression and secretion in vascular and immune cells.

In summary, the present study, for the first time, demonstrate that LXN plays a role in 1) SMC proliferation and migration through regulating PDGF receptor expressions and 2) macrophage migration though regulating ERK phosphorylation. While several factors have been implicated in mediating SMC proliferation and migration, PDGF is among the most vital phenotype-modulating factor in SMCs ^32^. Indeed, anti-PDGF or -PDGFR therapies have been shown to effectively inhibit neointimal hyperplasia ^32^. For example, the lack of PDGFR in SMCs has been shown to strikingly diminish neointimal hyperplasia after carotid artery ligation ^32^. Pharmacological inhibition of PDGF signaling reduces SMC proliferation and migration in the neointima ^32^. Besides, MCP-1/CCR2 also plays a crucial role in initiating atherosclerosis ^33^. Our present data strongly suggested that targeting LXN for inhibition may provide even more therapeutic benefits for treatment of cardiovascular disease such as neointimal hyperplasia and atherosclerosis through simultaneously inhibiting both SMC proliferation and macrophage infiltration.

## Non-standard Abbreviations and Acronyms

LXN: Latexin
SMCs: Smooth muscle cells
ECs: Endothelial cells
PDGF: Platelet-derived growth factor
PDGFR: Platelet-derived growth factor receptor
MCP-1: Monocyte chemoattractant protein-1
RNA: Ribonucleic acid
qPCR: quantitative real-time polymerase chain reaction

## Supplemental Materials

**Supplemental figure I.**
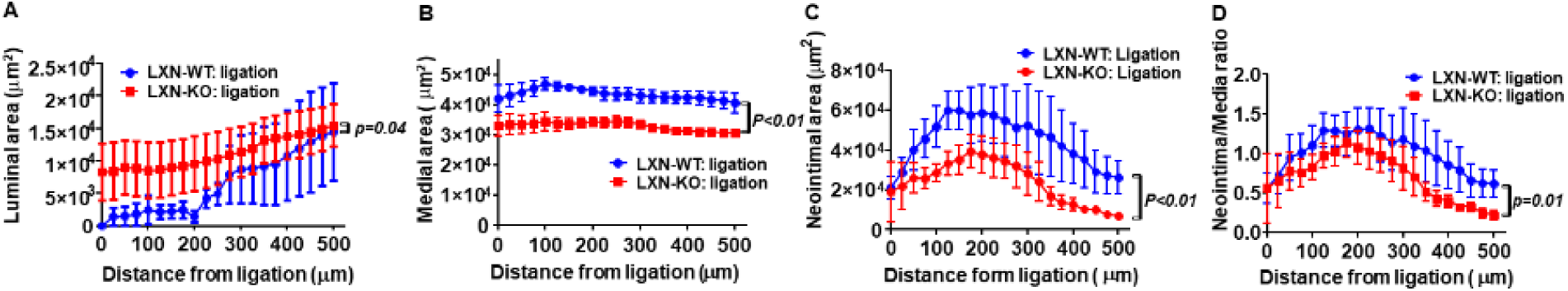
Histological analyses for neointimal hyperplasia with serial sectioning on LXN-gWT and -gKO. Global LXN knockout (LXN-gKO) or its control (LXN-gWT) mice were subjected to carotid artery ligation (left) or sham (right) surgery. 4-weeks after the surgery, carotid arteries were isolated and serial sections were cut. Luminal area (A), Medial area (B), Neointimal area (C), and Neointima/Media ratio (D) was measured by Image J software (n=4-5).

**Supplemental figure II.**
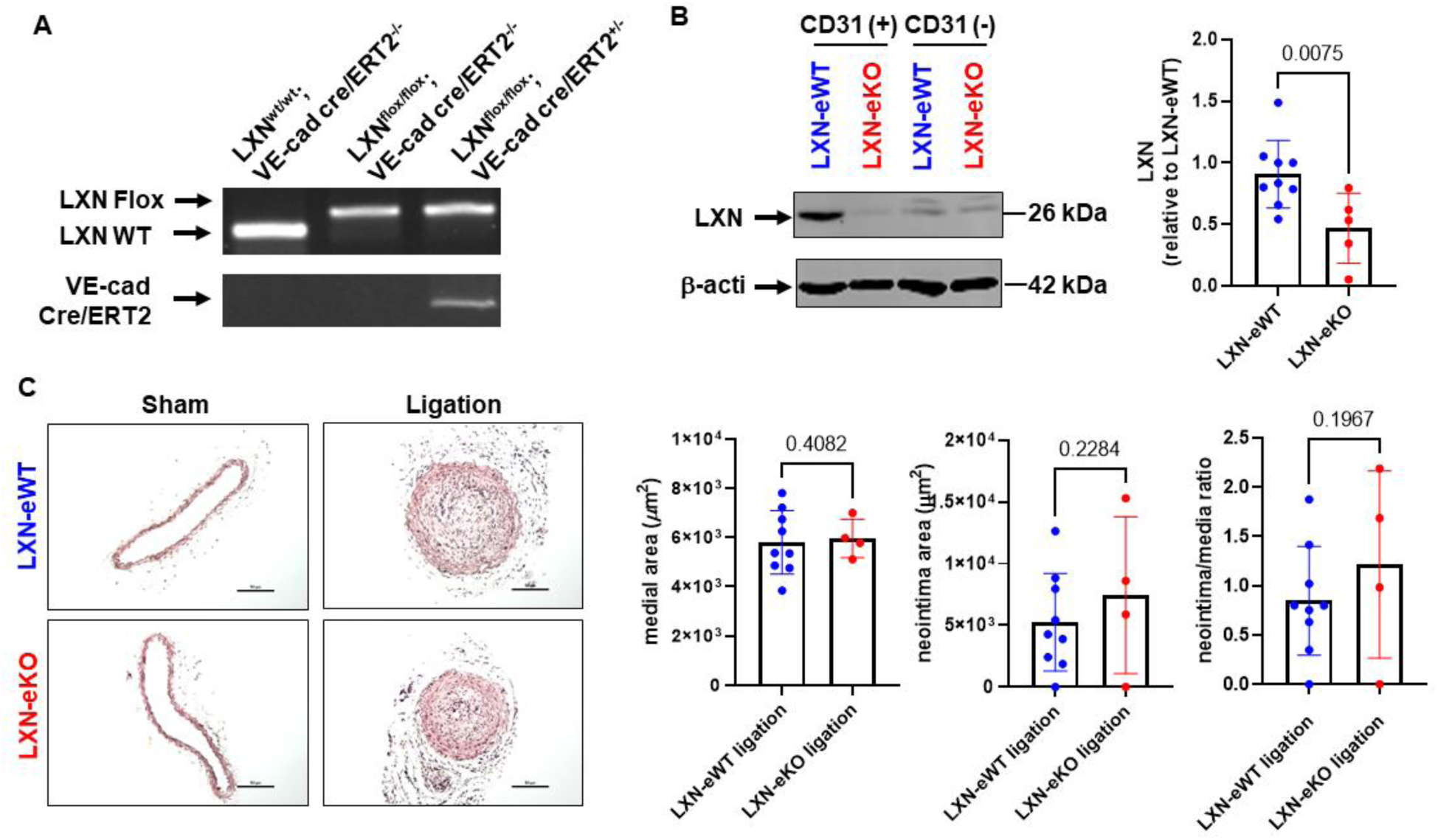
LXN-eKO does not suppress carotid artery ligation-induced neointimal hyperplasia. Inducible endothelial cell (EC)-specific LXN-KO (LXN^flox/flox^, VE-cadherin-Cre/ERT2 ^(+/–)^; LXN-eKO) and its control mice (LXN^flox/flox^, VE-cadherin-Cre/ERT2 ^(–/–)^; LXN-eWT) we used. A, Representative genotyping PCR images for LXN-flox and VE-cadherin cre/ERT2. 8-weeks old male LXN-eWT and -eKO mice were intraperitoneal injected with tamoxifen (75 mg/kg, 5 consecutive days) in corn oil. Mice were subjected to carotid artery ligation (left) or sham (right) surgery 1 week after the last tamoxifen injection at least. 4-weeks after the surgery, carotid arteries were isolated and used for histology. Simultaneously, mouse lung tissue were collected and primary mouse lung endothelial cells (MLECs) were isolated by magnetic-activated cell sorting with anti-CD31 antibody conjugated magnetic beads. B, Representative WB and quantification for LXN with MLECs isolated from LXN-eWT and -eKO. B-actin was used as internal control. C, Representative HE staining images. Scale bar=50 μm. Medial area, Neointimal area, and Neointima/Media ratio was measured by Image J software (n=4-9).

**Supplemental figure III.**
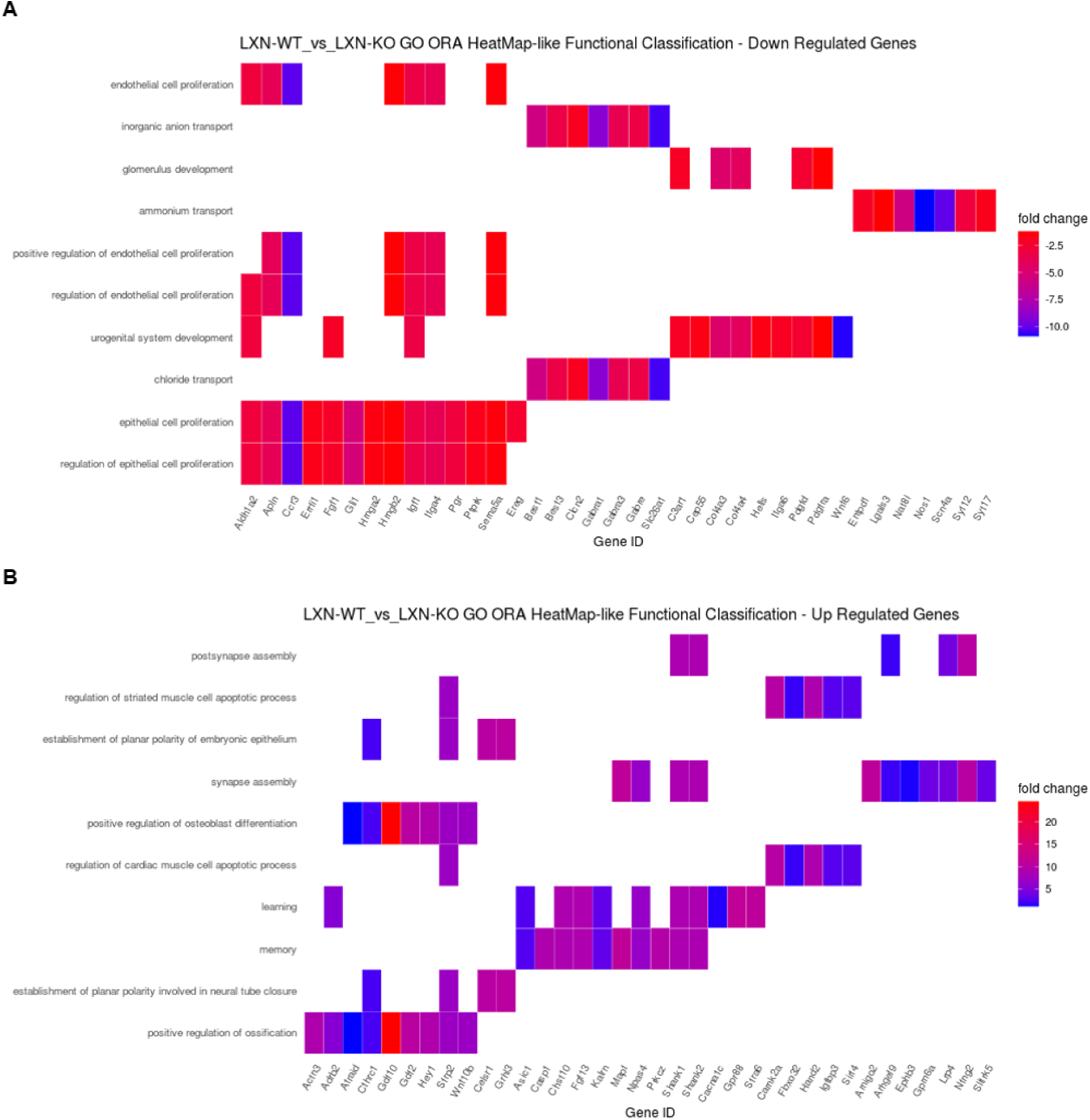
LXN deficiency suppresses proliferation-related genes in mouse SMCs. Primary mouse smooth muscle cells (SMCs) were isolated from LXN-gWT or -gKO aortas with the digestion method. Three aortas were used for one isolation. 2-independent primary SMC isolation was performed. Heatmap shows the functional classification of down-regulated (A) or up-regulated (B) genes by RNA-seq.

**Supplemental figure IV.**
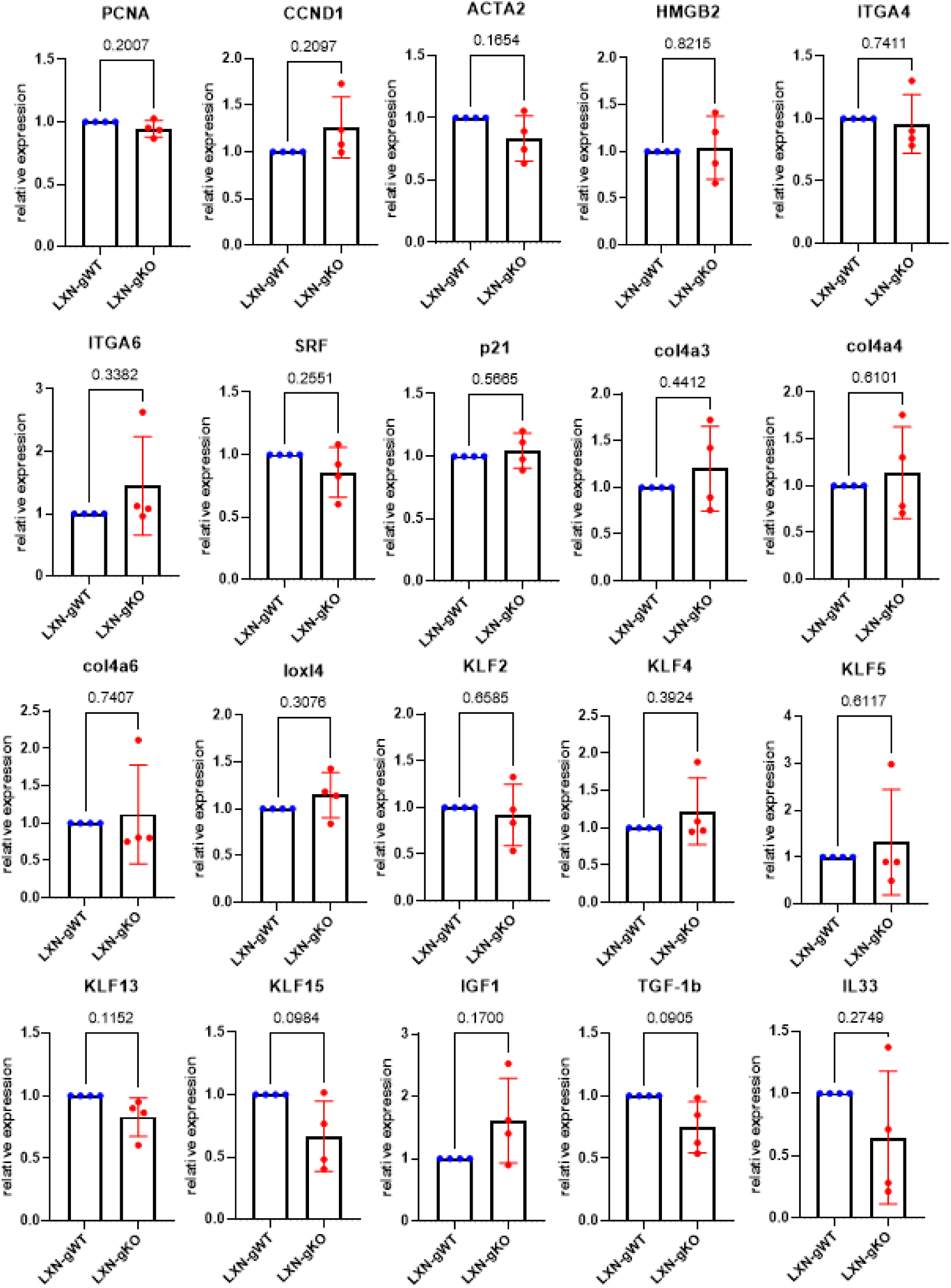
qPCR for SMC proliferation-related genes on mouse primary SMCs obtained from LXN-gWT or -gKO. Primary mouse SMCs were isolated from LXN-gWT or -gKO aortas with the digestion method. Three aortas were used for one isolation. Quantitative polymerase chain reaction (qPCR) was performed for potential LXN targeting SMC proliferation-related genes screened by RNA-seq.

**Supplemental figure V.**
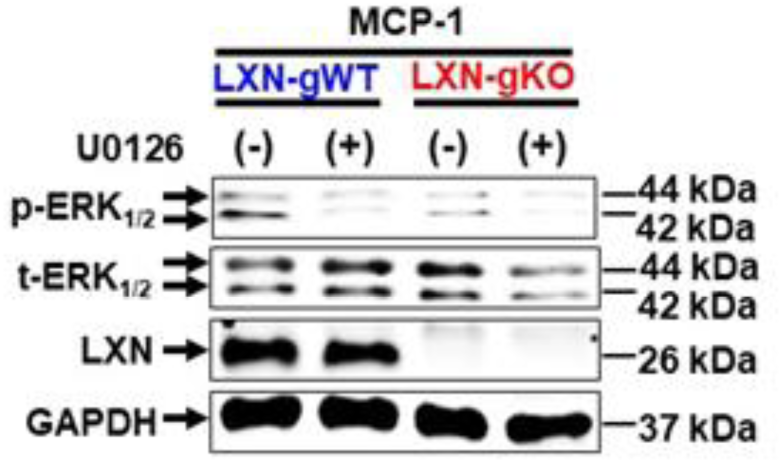
Western blot to confirm inhibitory efficiency of U0126. Representative WB for p-ERK, t-ERK, and LXN. PMs isolated from LXN-gWT or -gKO were stimulated with MCP-1 (100 ng/ml, 15 min) in the absence or presence of U0126 (10 μM, 24 h) (n=3). GAPDH was used as internal control.

**Supplemental figure VI.**
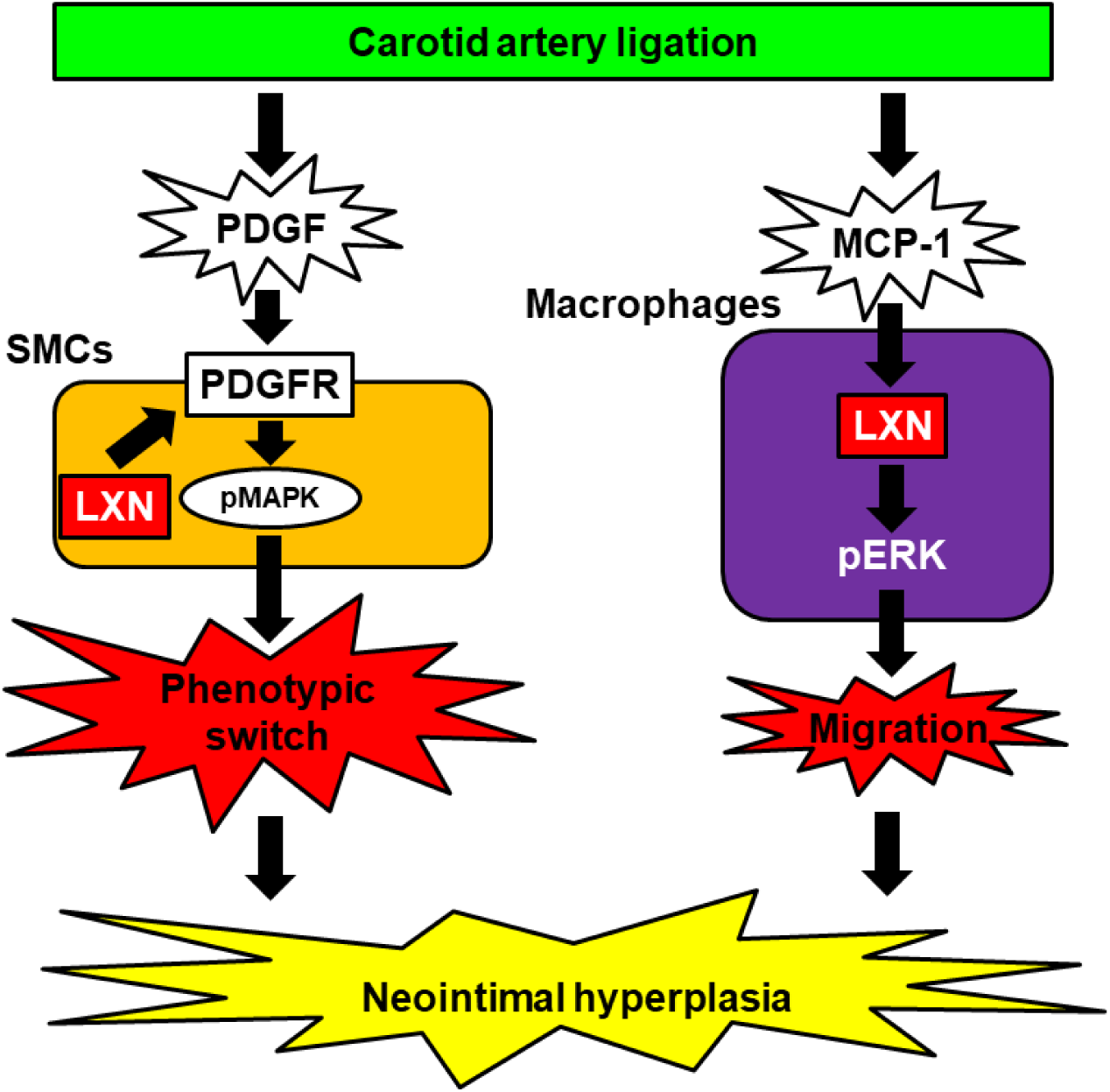
Schematic figure of summary.

